# Image-Based Quantitative Single-Cell Method Suggests Increase of Global Chromatin Accessibility in Tumor Compared to Non-tumor Cell Lines

**DOI:** 10.1101/2024.09.05.611456

**Authors:** Mairead Commane, Vidula Jadhav, Katerina Leonova, Brian Buckley, Henry Withers, Katerina Gurova

**Affiliations:** Department of Cell Stress Biology, Roswell Park Comprehensive Cancer Center, Elm and Carlton Str, Buffalo, NY, USA, 14263; Drug Discovery Core Shared Resource, Roswell Park Comprehensive Cancer Center, Elm and Carlton Str, Buffalo, NY, USA, 14263; Department of Bioinformatics and Biostatistics, Roswell Park Comprehensive Cancer Center, Elm and Carlton Str, Buffalo, NY, USA, 14263

## Abstract

The phenotypic plasticity of cancer cells has recently emerged as an important factor of treatment failure. The mechanisms of phenotypic plasticity are not fully understood. One of the hypotheses is that the degree of chromatin accessibility defines the easiness of cell transitions between different phenotypes. To test this, a method to compare overall chromatin accessibility between cells in a population or between cell populations is needed. We propose to measure the chromatin accessibility of a cell by total fluorescence signal from nuclei stained with DNA-binding fluorescent molecules. This method is based on the existing data that some small molecules bind nucleosome-free DNA more easily than nucleosomal DNA. Thus, nuclear fluorescence of these molecules is proportional to the amount of nucleosome-free DNA, serving as a measure of chromatin accessibility. We optimized the method using several DNA binding molecules and known chromatin modulating agents. Using a set of tumor and non-tumor cells of different origins we observed the tendency to the higher chromatin accessibility of tumor versus non-tumor cells. Chromatin accessibility was also increased upon oncogene-induced transformation of mouse and human cells.

## Introduction

Chromatin accessibility, or more precisely the accessibility of genomic DNA within chromatin for transcriptional machinery and other protein complexes, is a crucial determinant of a cell’s transcriptional program and phenotype. Cells vary in their ability to change phenotype, known as phenotypic plasticity, which plays a significant role in various physiological and pathological processes such as differentiation, oncogenic transformation, tumor progression, inflammation, etc [1]. However, we lack a reliable quantitative measure to assess and compare the degree of phenotypic plasticity between different cells.

Theoretically, phenotypic plasticity refers to a cell’s ability to switch between transcriptional programs. This can be measured by observing time-dependent changes in gene expression using techniques like bulk RNA-seq at different time points. While single-cell RNA sequencing (scRNA-seq) is a better approach for demonstrating the heterogeneity within a cell population, however, it is not sufficiently quantitative. Additionally, both methods are expensive, time-consuming, and require a high degree of technical expertise and sophisticated data analyses.

Current methods to measure chromatin accessibility as a potential proxy for phenotypic plasticity are extremely laborious and not considered to be high throughput. Techniques like ATAC-seq or nuclease-seq are designed to identify differences in accessibility between specific genomic regions but fail to quantify overall accessibility of chromatin of a cell to compare this parameter between cells [2]. The extensive biochemical and bioinformatic manipulations needed often raise questions about the reliability of these comparisons. Comparison of different samples requires the use of spike-in controls to normalize signals between different samples [3, 4] and not applicable to single-cell approaches. Image-based methods using confocal or superresolution microscopy are used to evaluate differences in chromatin condensation between different nuclear regions and compare staining patterns between cells [5–8]. These methods are excellent for studies of a specific chromatin-related research question; however, they still do not measure the total amount of accessible DNA per nucleus to compare this amount between cells in different states.

Therefore, a simple and reliable method to measure overall chromatin accessibility per cell is highly demanded. Extensive data indicates that the degree of chromatin condensation not only defines DNA accessibility to protein complexes, such as RNA polymerases, nucleases or transposases, but also to some small molecules [9–12]. The two most specific types of DNA ligands are DNA intercalators, heterocyclic compounds that insert their planar bodies between nucleoside bases of DNA, and minor groove binders, crescent-shaped molecules that position themselves inside and along the minor groove of DNA. Among both classes, there are many well-known fluorescent molecules, and significantly, the fluorescent signal of many of these molecules increases many folds when they are bound to DNA [13].

An interesting property found long ago among molecules within both classes, is that some of them bind free DNA more easily than DNA bound by proteins, especially histones within nucleosomes [9–12]. Thus, theoretically, the binding of these molecules to genomic DNA is proportional to DNA accessibility, and if they fluoresce only when bound, their fluorescence could be a measure of chromatin accessibility.

Several recent studies have proposed using DNA-binding small molecules to evaluate the chromatin state in cells (see Discussion) [6, 7, 14, 15]. Building on these previous studies, we propose and optimize an approach to measure the intensity of total nuclear fluorescence of DNA-bound small molecules as a reporter of chromatin accessibility in individual cells. Using several DNA intercalators, minor groove binders, and established methods to manipulate chromatin condensation, we provide evidence that the nuclear fluorescence of some of DNA ligands can serve as an easy and quantifiable proxy for chromatin accessibility in cells and tissues. Notably, this parameter is increased in cells with higher phenotypic plasticity, such as tumor and transformed cells.

## Results

### 1. Theoretical assumptions

There are studies showing that some DNA ligands bind preferably naked DNA versus DNA wrapped around nucleosomes [9–12]. Intercalator, such as propidium iodide or ethidium bromide bind DNA through the insertion of planar heterocyclic moiety between base pairs and this requires around a 2-fold increase of inter-base pair distance [16]. Spatial and superhelical constraints of nucleosomal DNA limit this binding [15]. Some histone’s amino acid side chains intercalate between base pairs, thus competing with DNA ligands. The minor groove of DNA facing histones is inaccessible to DNA minor groove ligands, such as DAPI or Hoechst. Reduced binding of some DNA ligands to nucleosomal versus nucleosome-free DNA was established in solution and cell-based experiments [9–12].

However, high-affinity DNA ligands compete with histones for binding of nucleosomal DNA. This causes DNA unwrapping from the histone core with loss of histones from chromatin, a phenomenon known as chromatin damage [17]. Therefore, such DNA ligands themselves cause an increase in chromatin accessibility, which makes measuring basal chromatin accessibility difficult. Another problem is that many DNA ligands are a substrate of multidrug transporters and their concentration in cells depends on the activity of multi-drug transporters, which is variable between cells [18]. Both these problems can be solved by cell fixation. The most suitable seems fixation with short distance crosslinking agents, since they cause covalent links between molecules, effectively gluing them together into an insoluble meshwork that preserves the molecular anatomy of a cell as it existed at the moment of fixation. Thus, no further nucleosome unwrapping is possible.

The next concern is the effect of the total amount of DNA in cells, which is different between G1, S, and G2 cells and may be different between healthy and diseased cells (e.g., normal and tumor cells due to aneuploidy, amplifications, and deletions). Thus, cells with longer total DNA would bind more DNA ligands without having more accessible chromatin. Therefore, normalization for the total DNA content per cell may be needed. However, the way to measure nuclear fluorescence may mitigate this issue.

To access overall chromatin accessibility per cell nucleus, total amount of DNA-bound dye needs to be quantified, what means collection of total fluorescent signal from the whole volume of a nucleus. Several recently published methods based on high-resolution confocal microscopy measured fluorescence from a section of a cell nucleus, not of the whole nucleus. Although images can be Z-stacked, they are still the sum of sections, and the total signal depends on the number and thickness of sections as well as thickness of a nucleus (Fig.1A). More accurate signal measurement from the whole nucleus is possible via using low resolution widefield microscopy with the focus depth of an objective (4X ∼ 32.5 microns, 10X ∼ 8.5 micrones) longer that nuclei thickness (5-10 micrones).

**Fig. 1.**
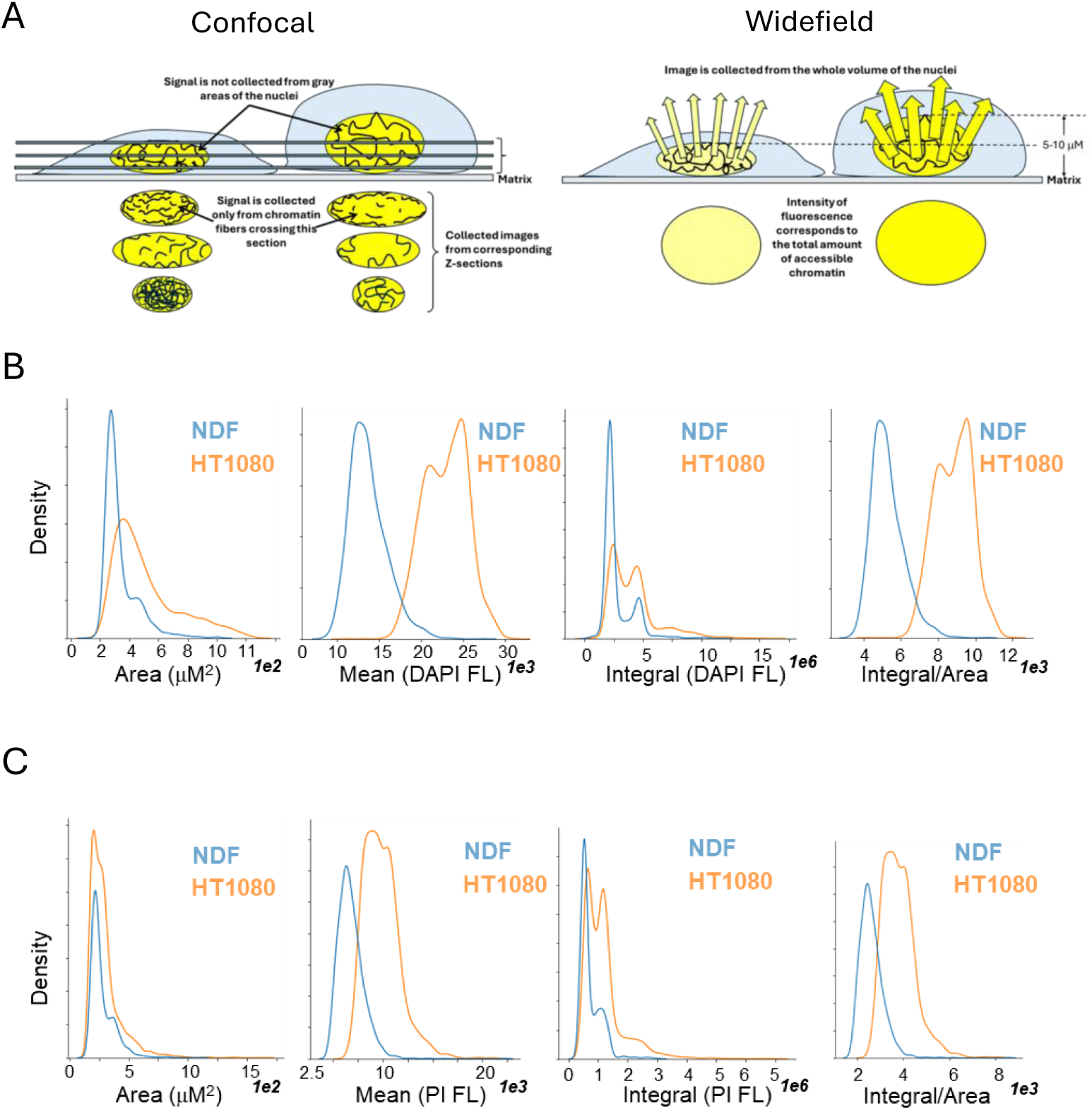
Measurement of nuclear fluorescence using non-confocal automated imaging. **A**. Total nuclear fluorescence is more accurately collected using non-confocal imaging. **B, C**. Distribution of parameters of nuclear fluorescence in NDF and HT1080 cells in basal conditions. Cells were fixed and stained with DAPI (**B**) or PI (**C**) 24 hours after plating.

Alternative methods are flow cytometry and non-confocal microscopy. Both methods allow separation of G1, S, and G2 cells, so the problem of cell cycle-dependent DNA content may be easily solved by assessing the cells in the same phase of the cell cycle, e.g., G1. However, flow cytometry has poor distinction between cytoplasm and nucleus (unless using special equipment like ImageStream) and ligand binding to mitochondrial DNA or double-stranded RNA in cytoplasm may affect the signal.

Automated widefield microscopy like flow cytometry allows collection of fluorescent signals from the whole volume of a cell (Fig.1A), however, the cytoplasm and the nucleus are easier separated by masking since cells are attached to the matrix. It also allows significant process miniaturization: as few as several cells can be detected in multi-well plates and it does not require cell detachment from a matrix on which they normally grow. It can also be applied to tissues without tissue disintegration (tissue slides). We tested effects of these theoretical considerations on the performance of a method in conditions of controlled chromatin decondensation in normal and tumor cells.

### 2. Optimization of an assay

#### 2.1. Selection of a parameter of nuclear fluorescence

We used normal human diploid fibroblasts NDF and human fibrosarcoma cells HT1080. The latter are near diploid (modal chromosome number = 46; range = 44 to 48) [19], thus eliminating the potential effect of cell aneuploidy. Cells growing in a multiwell plate were fixed and stained with intercalator propidium iodide (PI) or miner groove binder DAPI. Using widefield automatic image acquisition, we collected the following parameters: nuclear area, mean – the average intensity of all pixels per nucleus, and integral – the sum of all pixel intensities per nucleus. Images were collected using a 4X objective, with pixel size equal to ∼1.6 microns. These data are presented in the form of histograms in Figure 1B and C. As we expected, the area and the integral of cells had bimodal distributions reflecting the phase of a cell cycle similar to the distribution of cell fluorescence intensity measured by flow cytometry. Therefore, average integral values are significantly influence by the proportion of cycling cells in a population. However, the distribution of mean values did not follow the same bimodal pattern, which is explained by the cell-cycle dependent increase of the nuclear size. The normalization of integral by the area of a nucleus produced a distribution very similar to the mean (Fig.1B, C). Thus, we selected mean nuclear fluorescence as a potential measure of chromatin accessibility in cells due to less dependence on the phase of cell cycle.

#### 2.2. Comparison of DNA ligand

We tested the performance of several DNA ligands for which preferred binding to free DNA versus chromatin was already demonstrated, intercalator propidium iodide (PI), and minor groove binders DAPI and Hoechst [9], [20], [21], [22], as well an intercalator for which this was not well studied, SYBR Green. As an inducer of chromatin decondensation we used curaxin CBL0137 which destabilizing effect on nucleosomes was shown in different assays in solution and in cells [16, 23, 24]. We used a range of CBL0137 concentrations from 0.1 μM – no significant nucleosome unfolding, 0.3 μM – causing departure of H1 histone in some cells [25], 1-3 μM - loss of outer histones H2A and H2B, and 10 μM – significant nucleosome disassembly in cells ([26] and Fig. S1A). Importantly, these are approximate numbers, since these effects do not happen equally genome-wide but depend on the pre-existing chromatin state and cell type.

Cells were treated with CBL0137 for 10-60 min, since CBL0137 binds genomic DNA in cells in a matter of seconds [16]. Preliminary experiments done using flow cytometry showed that a shift in cell fluorescence is observed already after 10 minutes of incubation and is not significantly increased anymore between 20 minutes and 1 hour (Fig. S1B and C). After treatment cells were fixed and stained with PI, SYBR Green, Hoechst 33342 or DAPI. Imaging was done between 60 minutes and 24 hours with no difference between readings (data not shown).

The fluorescence of nuclei stained with different dyes increased with increasing doses of CBL0137 (Fig. 2A, B). Though the increase in average fluorescent signal per well was very small in case of minor groove binders (Fig.2A), it was highly significant when all individual cell values were counted (Fig.2B). The increase was stronger with intercalators, PI and SYBR Green.

**Figure 2.**
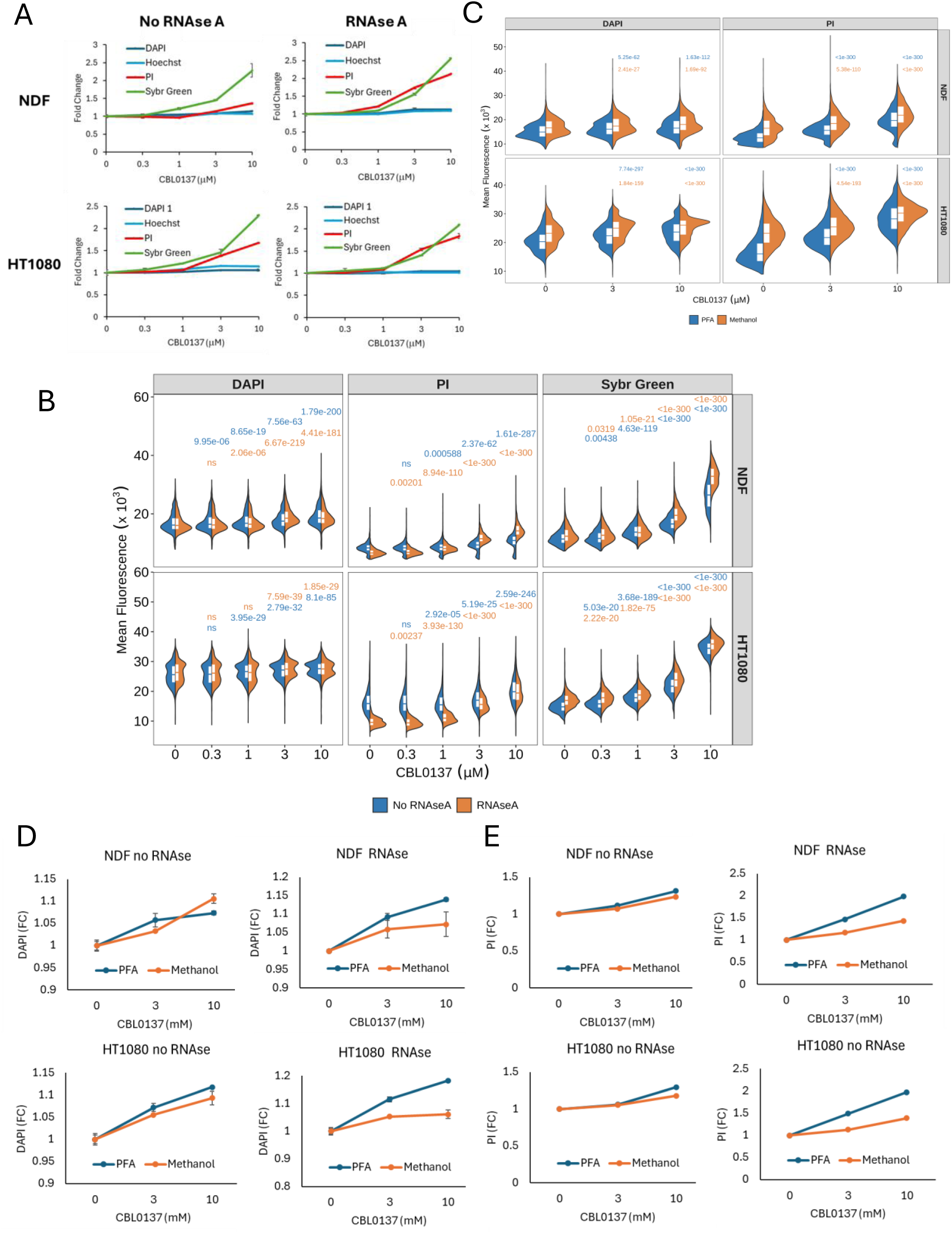
Optimization of the assessment of chromatin accessibility using fluorescent DNA ligands. Nuclear fluorescence was increased with a short-term treatment of NDF and HT1080 cells with nucleosome-destabilizing compound CBL0137. **A, B**. Comparison of the performance of different DNA dyes in the presence and absence of RNAse A. **A**. Fold change in average nuclear fluorescence per well relative to untreated cells. Data from two replicate wells with error bars showing variability between replicates. A representative from > 3 similar experiments. **B**. Split violin plots with quartiles showing mean nuclear fluorescence of cells treated with different concentrations of CBL0137 for 60 min. Numbers above violins are Holm adjusted p-values for the Kruskal-Wallis test with post-hoc Dunn’s test comparing treated cells to the untreated control stained either without RNAse A (blue numbers) or with RNAse A (orange numbers). P-values were rounded to 3 decimal places. ns - p-value >0.05. **C**. Split violin plots with quartiles showing mean nuclear fluorescence per cell fixed with either 4% PFA or 100% methanol. Numbers on the plot are Holm adjusted p-values for the Kruskal-Wallis test with post-hoc Dunn’s test comparing treated cells to untreated control fixed either with PFA (blue numbers) or methanol (orange numbers). **D, E**. Fold change in average nuclear fluorescence per well of cells treated with CBL0137 relative to untreated cells and stained in the presence of absence of RNAse A. **D**. Cells were stained with DAPI. **E**. Cells were stained with PI. Data from two replicate wells with error bars showing variability between wells.

CBL0137-induced increase of SYBR Green fluorescence was the strongest when cells were stained with 50 μM of SYBR Green. However, when we titrated the concentration of dyes in staining solution, PI fluorescent intensity decreased linearly with dye concentration, while SYBR Green fluorescent intensity was bell-shaped (Fig. S2A, B). We confirmed this non-linear change in fluorescence by inspection of images taken with different exposures (Fig. S2 C, D). For both intercalators, the CBL0137-induced increase of fluorescent signal became smaller with increase in the intensity of fluorescence in basal conditions, suggesting a saturation effect, which was especially strong in the case of SYBR Green (Fig. S2E, F). With this dye automatic quantitation without visual confirmation may be misleading due to easy overexposure (compare visual effect and quantitation in Fig. S2 D and F).

#### 2.3. Unwrapping versus untwisting

Binding CBL0137 to DNA causes not only unwrapping of DNA from the nucleosome core, but also untwisting of DNA helix (Fig. S3A). The question was whether helix untwisting would affect DNA dyes binding and fluorescence, thus interfering with the assessment of DNA unwrapping. The test the role of helix untwisting in DNA dye fluorescence we used a linearized DNA plasmid and incubated it with increasing concentrations of CBL0137. After that the sample was run in agarose gel. We also run in parallel supercoiled plasmid incubated with CBL0137 which allows easier observing the shift of supercoiled DNA due to CBL0137 intercalation. Gel was stained either with PI or Hoechst after running (Fig.S3B). Fluorescence of both dyes was measured using band intensity tools in Image Lab Software (BioRad). Untwisting of linear DNA helix caused by CBL0137 did not change PI fluorescence in any of experiments and replicates (Fig.S3C), while Hoechst fluorescence was minimally reduced (within 1-2 %, Fig. S3D, E), reaching statistical significance only at the highest concentration of CBL0137. Thus, we concluded that DNA helix untwisting did not increase dye binding to DNA

#### 2.4. Importance of RNAse A treatment

Although both intercalators and minor groove binders are known as specific DNA dyes, they also bind to double-stranded RNA (dsRNA). Thus, we compared the performance of all DNA dyes with and without treatment of fixed cells with RNAse A. Distribution of fluorescent signal was changed upon RNAse A treatment in all conditions (Fig. 2B – compare blue and orange halves of volcano plots), however the most significant change was seen for intercalators, especially for PI (Fig. 2A, B and Fig. S4). This change was explained by stronger cytoplasmic signal in cells stained with intercalators in the absence of RNAse A (Fig. S4). Cytoplasmic signal was gone upon RNAse A treatment, suggesting that intercalators stain dsRNA stronger than minor groove binders. Removal of RNA staining with RNAse A treatment made changes in chromatin accessibility caused by CBL0137 more pronounced.

#### 2.5. The role of fixation methods

Although we assumed that fixation with the short-range crosslinking agent would be the best, we decided to see if other fixatives, such as methanol, which causes protein denaturation, can be used for the same purposes. Methanol fixation led to an increase in fluorescent signal in all conditions compared with PFA fixed cells (Fig. 2C). There was a similar increase in DAPI and PI nuclear fluorescence upon CBL0137 treatment if cells were not treated with RNAse A. However, RNAse A treatment led to a further increase of the effect of CBL0137 if cells were fixed with PFA and no additional increase if cells were fixed with methanol (Fig. 2D, E). The latter may be explained by a higher level of RNA degradation from methanol fixation than from PFA, especially since we used PBS for rehydration [27].

Thus, we defined the following conditions as the most sensitive and robust for detection of changes in chromatin accessibility: (i) fixation of cells with 4% PFA, (ii) treatment of cells with RNAse A, (iii) staining of cells with DNA intercalator dyes, with PI being less sensitive to changes in dye concertation but more dependent on RNAse A treatment, and SYBR Green requiring more accurate concentration optimization and monitoring of exposure time, due to easy overexposure.

### 3. Testing of the assay performance in cells treated with epigenetic drugs

After establishing optimal conditions, we tested assay performance upon treatment of cells with several epigenetic drugs with known effects on chromatin accessibility. We used HDAC inhibitors, trichostatatin A (TSA), panobinostat (PNB), valproic acid (VA) and vorinostat (SAHA) and inhibitor of BET domain proteins, JQ1, which does not directly increase chromatin accessibility. Since most of these drugs inhibit enzymes, we treated cells for 24 hours and monitored the toxicity by direct cell counting and microscopic imaging (Fig. 3A, B, E, F). As expected, cells treated with all HDAC inhibitors accumulated more PI than untreated cells (Fig. 3C-F), in line with the known mechanism of action of these agents. This effect was obvious and significant even when the drug caused growth arrest (reduced number of cells per well compared with control in the absence of cell death) in the case of PNB and SAHA, or cell death in the case of VA (Fig. 3A, B, E, F). Interestingly, all these drugs had a stronger effect on the viability and chromatin accessibility of tumor HT1080 cells, than normal NDF cells (Fig. 3). JQ1 caused the minimal changes in chromatin accessibility (Fig. 3 C, D) in line with its mechanism of action.

**Figure 3.**
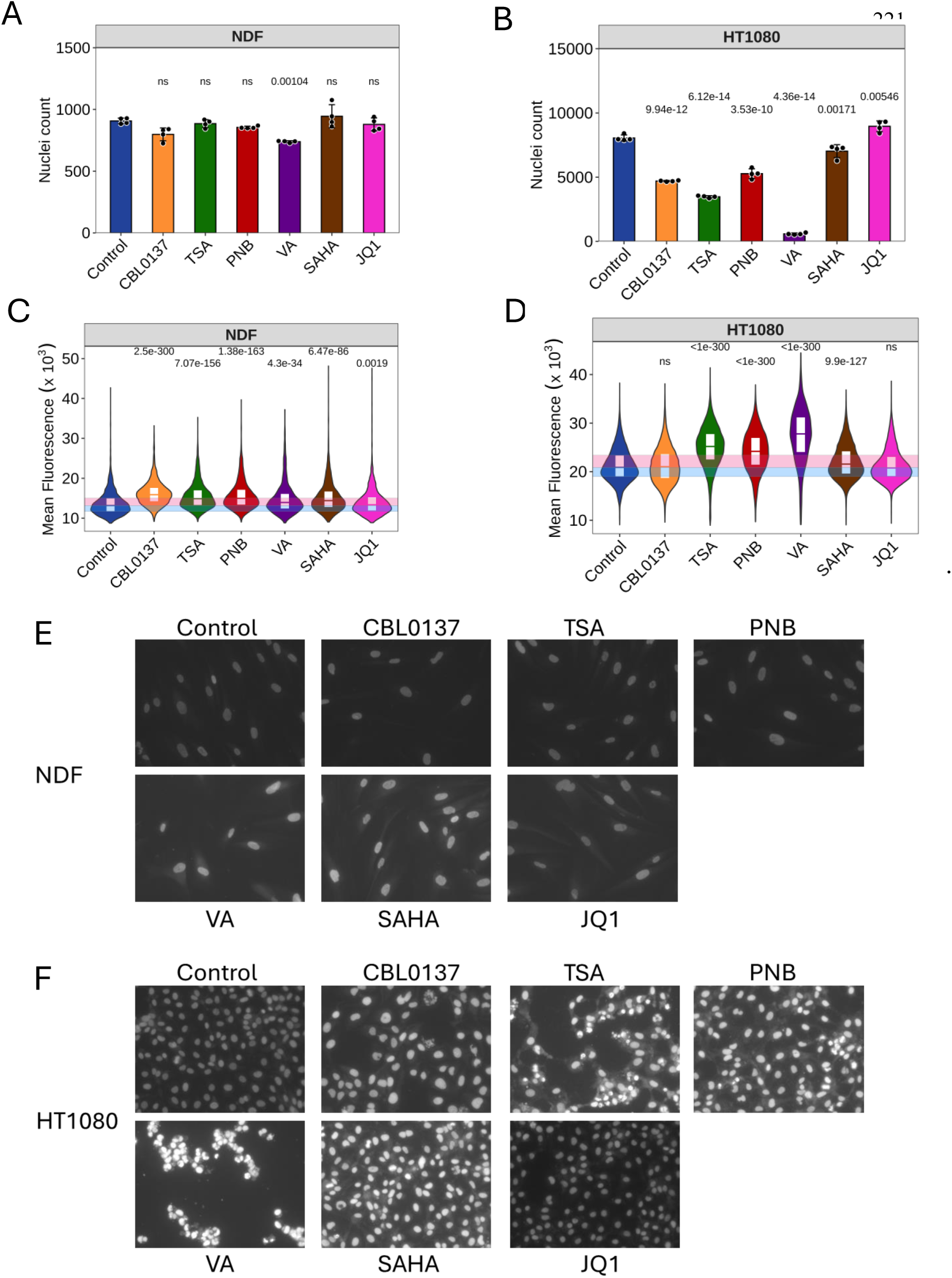
Effect of epigenetic drugs on chromatin accessibility measured with PI. NDF and HT 1080 cells were treated with CBL0137 (500 nM), trichostatin A (TSA, 500 nM), panobinostat (PNB, 20 nM), valproic acid (VA, 0.5 mM), SAHA (1000 nM), or JQ1 (1000 nM) for 24 hours before fixation with PFA and staining with PI in the presence of RNAse A. **A, B**. Number of nuclei per well of NDF (**A**) or HT1080 (**B**) cells, untreated or treated with the drugs. Bars are mean value of 4 replicate wells, error bars are SDV. Number above the bar is p-value for ANOVA with post-hoc Tukey test comparing each drug with the control untreated cells, rounded to 2 decimal points. **C, D**. Violin plots with quartiles of distribution of PI nuclear fluorescence of cells in all replicate wells of NDF (**C**) or HT1080 (**D**) cells treated with epigenetic drugs. Number above violin is Holm adjusted p-value for Kruskal-Wallis with post-hoc Dunn’s test comparing each drug with the control untreated cells, rounded to 3 decimal points. . **E, F**. Microscopic images of PI-stained wells from the plate of cells treated with epigenetic drugs. The same exposition time was used for all images of NDF (**E**) and HT1080 (**F**) cells.

To test if our method would report decrease of chromatin accessibility in cells, we used three established methods to cause chromatin compaction in HT1080 cells: hypertonic shock via 60 minutes cell incubation in 300 mM NaCl solution, acute depletion of ATP via 30 min cell incubation in 10 mM sodium azide 50 mM 2-deoxyglucose in PBS, and treatment of cells overnight with histone demethylases inhibitor methylstat [28]. The latter is an inhibitor of Jumonji C domain-containing histone demethylases, which include demethylases erasing inhibitory and activating histone modifications H3K9me, H3K4me, H3K36me, H3K27me, and several others. However, due to a much higher presence of inhibitory H3K9me3 mark in the mammalian genome, the demonstrated effect of methylstat in chromatin compaction [29]. All three treatments led to the decreased staining of nuclei, both PI and DAPI (Fig. S5).

Thus, our method detected chromatin accessibility changes caused by several groups of epigenetic drugs though neither of these drugs bind DNA.

### 4. Change of chromatin accessibility during cell cycle

The amount of DNA-bound small molecules increases with DNA length. Since DNA length increases with cell cycle progression populations of cells with more cycling cells would bind more DNA dye due to longer average DNA length per cell. We tried to distinguish the effect of increased DNA length from increased DNA accessibility by comparison of PI incorporation at different phases of the cell cycle and in growth-arrested and cycling cell populations.

First, we identified cells in the S phase by labeling them with EdU for 1 hour and then measuring the fluorescent signal of DNA ligand in cells positive and negative for EdU. This comparison showed that EdU-positive cells accumulate more dye molecules, which was not surprising since they have more DNA than cells in G1, the predominant state of EdU-negative cells (∼ 70 % of HT1080 cells in basal conditions) (Fig. 4A, B). To mitigate this factor to a certain extent, we used the integral of cell fluorescence to identify cells in G1 or G2/M phases of cell cycle (Fig. 4B). EdU positive cells withing these groups are cells which are early in S phase and therefore having DNA content close to G1 cells or very late in S phase, i.e., having DNA content close to G2/M cells. When we compared mean fluorescent signal of these cells, we did not see significant difference between EdU positive and negative cells (Fig. 4C, D), suggesting that DNA replication per se does not make chromatin more accessible at nuclei level.

**Figure 4.**
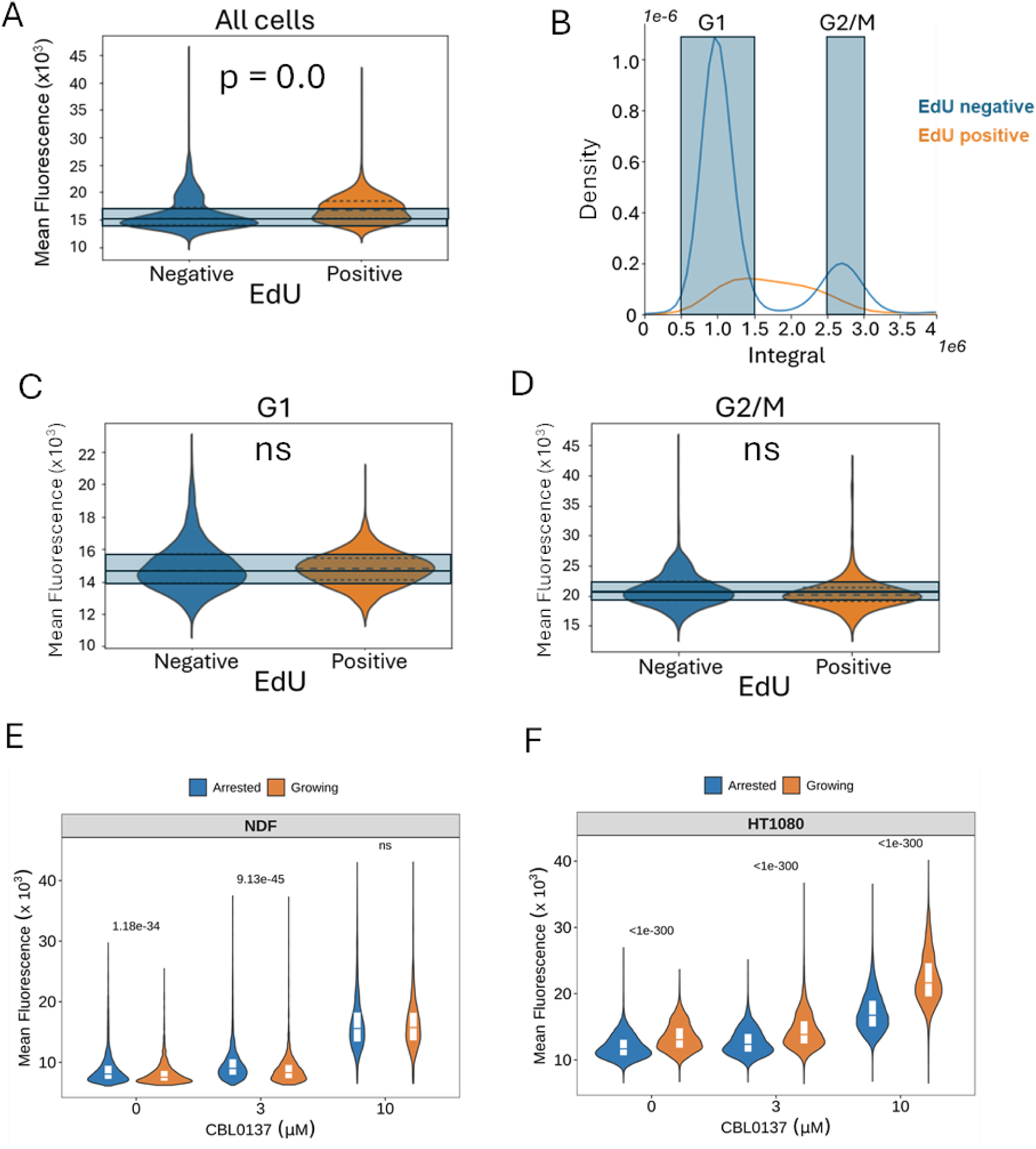
Dependence of chromatin accessibility of cell proliferation. **A-D**. HT1080 cells were incubated in the presence of EdU for 1 hour. After that cells were fixed and stained for EdU and DAPI. **A**. Violin plots with quartiles of nuclear fluorescence of of EdU positive and negative cells. Bleu squares correspond to the positions of quartiles (0.25, 0.5 and 0.75) in EdU negative cells. **B**. Histogram of distribution of total nuclear fluorescence (Integral) in EdU negative and positive cells. Blue squares show values of nuclear fluorescence used to select cells with DNA content close to G1 or G2/M cells. **C**, D. Violin plots with quartiles showing mean nuclear fluorescence of EdU positive and negative cells with DNA content close to G1 (**C**) or G2 /M (**D**). ns – p-value> 0.05. **E, F**. Comparison of mean nuclear fluorescence of NDF (**E**) or HT1080 (**F**) cells growing in normal conditions or arrested with dense plating and medium with 1% FBS for 48 hours. Before staining cells were treated with the indicated concentrations of CBL0137 for 60 minutes. Numbers are Holm adjusted p-values for Kruskal-Wallis with post-hoc Dunn’s test comparing growing and arrested cells at each concentration of CBL0137.

Another approach was to compare cells arrested with combination of contact inhibition and serum starvation (48 hours in medium with 1% FBS) and normally growing cycling cells. Interestingly, there was or slight decrease in basal condition and no difference upon CBL0137 treatment in the dye accumulation in growing versus arrested normal NDF cells, while growing tumor HT1080 cells had more open chromatin. This shift was true even if we looked only at the position of G1 cells using the integral of fluorescence, and it was increased at high dose of CBL0137 (Fig. S6).

Thus, we observed more open chromatin state in cycling versus resting cells, especially in the case of tumor cells and this increase cannot be explained by increased DNA length.

### 5. Comparison of chromatin accessibility in normal and transformed or tumor cells

Existing data from literature suggest that chromatin may be overall more open in tumor than in normal cells (reviewed in [30]), though data comparing general chromatin accessibility between normal, transformed, and tumor cells are missing. Thus, we decided to test this using our method.

First, we compared NDF and HT1080 cells using DAPI and PI. In both cases HT1080 accumulated more dye than NDF cells in basal conditions and upon CBL0137 treatment (Fig.5A, B). This was true with both fixatives, PFA and methanol, and in the presence or absence of RNAse A (Fig. S7A-D). Next, we tested two additional sets of tumor/non-tumor cells of breast and kidney origin. The first set included immortalized mammary epithelial cell line MCF10A and three breast cancer cell lines of different types, including triple negative MDA-MB-231 and BT549 cells and hormone receptor positive MCF7 cells. Kidney set consisted of spontaneously immortalized kidney epithelial cells and three renal cell carcinoma cell lines, ACHN, RCC45, and RCC54 [31]. A higher mean nuclear fluorescent signal was observed in tumor cells versus non-tumor cells in both sets (Fig.S8C-F). Although all these cells are non-syngeneic, and some of them have a ploidy different from 2n, their nuclear size changes in line with ploidy (Fig. S8A,B), partially normalizing the difference in DNA content when comparing the mean fluorescence of a nuclear pixel.

**Figure 5.**
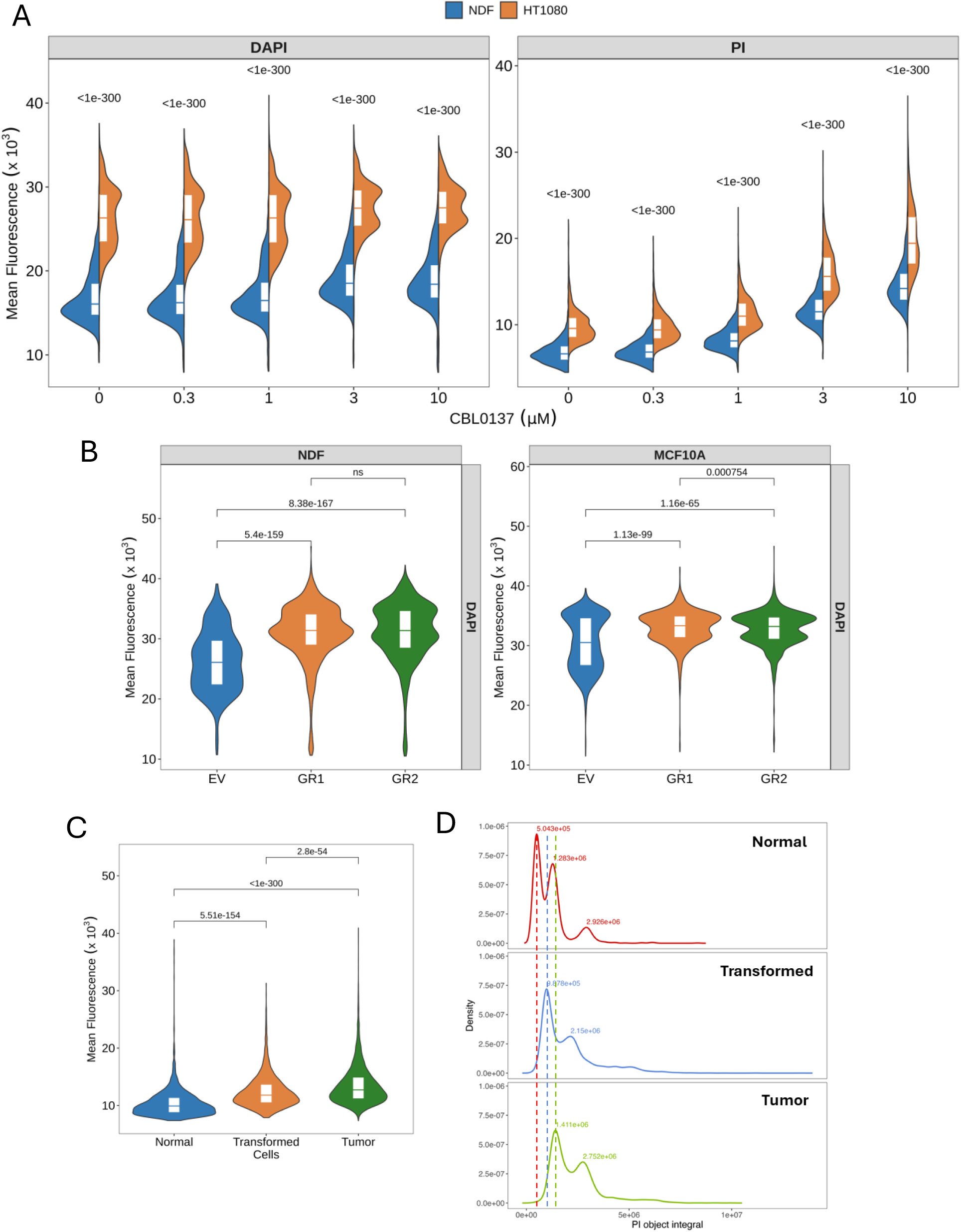
Comparison of chromatin accessibility between normal, transformed and tumor cells. **A, B**. Split violin plots of mean nuclear fluorescence of NDF and HT1080 cells treated with different concentrations of CBL0137 for 60 minutes and stained with DAPI (**A**) or PI (**B**). Numbers are Holm adjusted p-value for Kruskal-Wallis with post-hoc Dunn’s test comparing NDF and HT1080 cells. **C, D**. Violin plots with quartiles showing mean nuclear fluorescence (DAPI) of two biological replicates of transformed NDF (**C**) or MCF10A (**D**) cells (GR1, GR2) and control non transformed cells (EV). Numbers are Holm adjusted p-values for Kruskal-Wallis with post-hoc Dunn’s test comparing each of transformed variants (GR) with control non-transformed EV cells. **E, F**. Comparison of nuclear fluorescence (PI) of mouse normal, transformed and tumor cells. **E**. Violin plots with quartiles of mean nuclear fluorescence. Numbers above the lines show Holm adjusted p-values for Kruskal-Wallis with post-hoc Dunn’s test comparing all groups. **F**. Distribution of total nuclear fluorescence. Numbers show X-values of corresponding peaks on histograms. Dotted lines correspond to the positions of G1 peaks of normal MEF (red), transformed (blue) and tumor (green) cells.

Next, we compared syngeneic normal and transformed cells. NDF and MCF10A cells were transduced with combination of p53 dominant negative inhibitor, GSE56 [32] and mutant HRasV12 oncogene virus (GR) or empty virus (EV). Cells were passaged for 2 weeks to allow transformation phenotype to evolve and then they were fixed and stained with DAPI. Though nuclear accumulation of this dye has been changed less upon CBL0137 treatment than that of PI, we observed a significant difference between cells transduced with the EV and two independently generated transformed cell lines (GR1 and GR2). Both, transformed fibroblasts and epithelial cells accumulated more dye than non-transformed cells (Fig. 5C, D), suggesting more open chromatin state in transformed versus normal cells.

Lastly, we compared chromatin accessibility of syngeneic mouse normal, transformed and tumor cells. We used mouse embryo fibroblasts from wild type and p53 knockout animals. The latter were transduced with mutant HRasV12 to get transformed cells. These transformed cells were implanted subcutaneously into C57Black/6 mice and after tumor reached the size of 500 mm^3^, it was excised, disaggregated and established as cell line *in vitro*. Similar to human cells, we saw increase in chromatin accessibility upon *in vitro* transformation of mouse cells and further increase in tumor cells (Fig. 5E).

Since, as we saw in the previous section, mean fluorescence is higher in proliferating than non-proliferating cells, we decided to confirm that the increase in chromatin accessibility is not only due to the higher proportion of proliferating cells in transformed and tumor cell populations. For this, we compared fluorescent intensities of G1 peaks obtained from the integral of fluorescence. In all cases we saw shift in the position of G1 peaks corresponding to the shift of mean fluorescence (Fig. 5F and S5 E-H), confirming increase in the chromatin accessibility during oncogene induced transformation of human and mouse cells.

## Discussion

The method we described here is extremely simple and was used extensively for a different purpose: to assess the distribution of cells in populations along different phases of a cell cycle via flow cytometry. In most cases, laser settings were adjusted for every different cell type, and therefore, shifts in the positions of histogram peaks between cell types, i.e., different fluorescent intensities of different cells, were largely ignored by cell and molecular biologists. At the same time, the fact that PI or DAPI bind naked DNA better than DNA wrapped around histone core was observed long ago [9–12]. Interestingly, for PI it was also shown that it can bind to histones; however, while binding to DNA significantly increases PI fluorescence, binding to histones further reduced weak fluoresce of free PI [33]. In line with this, Bosire et al. showed that PI intensity negatively correlates with the anumberof histones in cells suggesting that major signal comes from PI binding to DNA [15].

The differential accessibility of DNA in chromatin to different molecules is currently mostly assessed using proteins, such as nucleases or transposases. These methods are not quantitative enough at the single-cell level, they are also expensive, time-consuming, and require high-level expertise in molecular biology and bioinformatics for data processing. Most importantly, they are not tuned for comparing the total level of accessible chromatin between individual cells and between different samples.

Nucleosome-dependent binding of small molecules to DNA in cells manifested with the emission of light presents an easy opportunity to assess an abundance of nucleosome-free DNA in different cells by comparing their nuclear fluorescence. This comparison can be made between cell populations and between individual cells in the population. Not surprisingly, there were already several attempts to use fluorescent DNA ligands for the assessment of chromatin organization in cells. The closest approach was published by Rosevalentine Bosire et al [15]. They used different methods to decondense chromatin in fixed cells to demonstrate that intercalator incorporation into nuclear DNA in cells is constrained by superhelical stress due to DNA wrapping within nucleosomes. Unlike us, they used laser scanning cytometry, i.e., they collected the signal from a section of a nucleus and not from the whole nucleus. They observed that the fluorescent signal of intercalators depends on the degree of chromatin condensation and negatively correlates with the number of histones per nucleus. Contrary to us, they did not see the same dependence for minor groove binder DAPI when they fixed cells with methanol, probably due to the assessment of a signal from one section of a nucleus.

Differential binding of an intercalator to open and closed chromatin was used by Gali Bai et al to develop a different method, adduct sequencing (Add-seq), to probe chromatin accessibility by treating chromatin with the small molecule angelicin, which then was covalently bound to DNA. DNA regions with bound angelicin was detected by nanopore sequencing [14]. Though Add-seq is similar to our method based on small molecule binding to DNA, this method requires nuclei isolation, DNA sequencing, and it is unclear whether it is good for measurement of differences in general chromatin compaction between cells and whether it can be applied to single cells.

Application of DNA ligands to discern chromatin structure between tumor and normal cells was proposed by Jianquan Xu et al [6]. They used minor groove binder Hoechst and stochastic optical reconstruction microscopy (STORM) [34] to evaluate chromatin organization. However, instead of Hoechst fluorescence, which is not optimal for super-resolution, they attached another fluorophore, Cy5, to Hoechst. Cy5 does not bind DNA itself but provides necessary fluorescent parameters for STORM. They observed significant differences between tumor and non-tumor cells, but their approach to image analysis was focused on the detection of different patterns of chromatin organization and not on the total cumulative fluorescent signal from a cell [6]. However, their description of the observed differences was in line with the quantitative differences that we observed here. They found that in normal cells chromatin domains are more compact, especially at the nuclear periphery. In precancerous cells, chromatin compaction was slightly disrupted, and in cancer cells, chromatin at the nuclear periphery was indistinguishable from the interior. This can be interpreted as general chromatin de-compaction in the process of tumorigenesis. This is an excellent method to detect and describe changes in the patterns of chromatin organization. However, it is much more complicated in data acquisition and analysis than the method we propose. Our method is simple enough that it can be done in a clinical lab. With additional optimization it can be applied to tissue slides, and data processing can be easily automated. We believe that the degree of chromatin decondensation may be an important measure of tumor aggressiveness [30]. Therefore, quantitative assessment of the chromatin decondensation in tumor cells and/or the proportion of cells in tumors with decondensed chromatin may be used as a prognostic marker.

Metastasis, invasion to the neighboring organs, and development of resistance to multiple therapies are factors responsible for poor prognosis of cancer patients. These traits are not often due to the accumulation of new mutations and selection of resistant clones, but to the adaptation of tumor cells to new conditions via changing of their phenotype, i.e., phenotypic plasticity [1, 35]. Although detailed mechanisms of phenotypic plasticity are still obscure, easiness of transitions between transcriptional programs is probably an obligatory factor of phenotypic plasticity. Transcriptional programs are controlled by chromatin organization at different genomic regions and therefore stable programs should be associated with stable chromatin with multiple constraints preventing easy activation and de-activation of genes. Easy transitions between active and inactive state of transcription may occur when silent and active states of genomic regions are not enforced strongly enough by chromatin organization, which may include unstable nucleosomes at regulatory region and gene bodies leading to easier access of transcriptional machinery to DNA, mobile, flexible chromatin fibers and “poorly locked” chromatin loops making random collisions of promoters and enhancer more probable and burst of transcription more stochastic. Thus, our long-term hypothesis was that general destabilization of chromatin through reduced number of histones per cell, prevalence of histone modifications or mutations making nucleosomes less stable, loss of heterochromatin proteins or overexpression of proteins making nucleosomes more open, e.g., HMG proteins, these all would result in more open chromatin state in tumor versus non-tumor cells. Our quantitative method confirmed this hypothesis in human and mouse models. Our next hypothesis is that the degree of chromatin decondensation correlates with tumor prognosis can now be tested by staining of patient’s tumor samples on slides and assessing correlation between the degree of chromatin accessibility and patient outcomes. If proven correct, this may become one of the simplest and universal prognostic biomarkers, which in case of development of appropriate fluorescent probe can be even applied for *in vivo* imaging.

## Limitations of the study

Although we believe that our method is simple, easy, quantitative, and accurately reveals the state of chromatin in cell populations and the difference in chromatin condensation between individual cells in a population, there are still limitations or uncertainties that need to be clarified in future studies. Ideally normalization for the total DNA length needs to be done. Thus, in the present state, this method is good for experimental studies, though the clinical application will require measurement of the total DNA content per cell. This might be achieved by decrosslinking of a sample followed by full unwrapping of DNA from nucleosomes with agents such as curaxin and another round of staining with fluorescent DNA ligand. A better understanding of the mechanisms of fluorescence as a result of interactions of molecules of DNA ligand between themselves and with DNA is needed to understand which signals report accessible chromatin and which are influenced by ligand-ligand interactions, as we see in the case of SYBR Green. Another question is which DNA ligand better reflects the actual chromatin state in different conditions, minor groove binders, or DNA intercalators. There is very limited data to understand which properties of nucleosomal DNA define these molecules binding. Most probably, their binding would inform us about the different complementary properties of nucleosomes. which can be put together to better understand chromatin organization and dynamics. Although fluorescent signals from the same cells in different experiments were in the same range, there were some differences between experiments and between wells in the same experiments. For comparison of different samples, especially in clinical conditions, a set of controls for normalization of staining and imaging and calibration of signal need to be developed.

## Acknowledgments

We would like to acknowledge Dr. Razvan Chereji for helping with Fig. S1A presentation, Drs. Andrei Gudkov and Subhamoy Dasgupta for the critical discussion of the manuscript, Bruce Specht, Mary Morgan, Sarah Mercy and Dale Henry for the administrative support.

## Funding

National Institutes of Health grant R01 CA266216 (KG) National Institute of Health grant NCI NIH Core P30CA16056 to RPCCC, which partially covers the costs Genomics, Bioinformatics, Flow Cytometry, Drug Discovery Lab, Gene Modulation shared resources.

## Author contributions

MC and VJ were responsible for running imaging and flow cytometry experiments. KL generated mouse transformed and tumor cells. BB did imaging and image analyses, HW performed analysis of all data, generated R script to define x-value of positions of G1 and G2 peaks on a histogram and editing of the manuscript. KG conceptualized the study, analyzed all data, wrote and edited the manuscript as well as prepared some figures.

## Competing interests

Authors declare that they have no competing interests.

## Data and materials availability

Transformed or tumor cells generated for this study are available upon request. Raw imaging data are available upon request. R script to define x-value of positions of G1 and G2 peaks on a histogram is available here: https://github.com/HGWithers/cellphaseR

## Materials and Methods

### Reagents

CBL0137 was provided by Incuron, Inc (Buffalo, NY). Propidium Iodide, RNAse A, Trichostatin A, Panobinostat, Valproic Acid, Vorinostat (SAHA), JQ1, methylstat, sodium azide and 2-deoxyglucose were purchased from Sigma-Aldrich (St. Louis, MO), DAPI, Hoechst 33342 and SYBR Green were purchased from Invitrogen/TermoFisher (Grand Island, NY). EdU kit was from Click Chemistry Tools (Scottsdale, AZ).

### Cells

HT1080 (male) and MCF10A, MCF7, MDA-MB-231, BT549 (female) cells are from the American Type Culture Collection (ATCC). Ht1080 and MCF7 cells were authenticated using short tandem repeat analysis (100% match). Primary human neonatal dermal fibroblasts (NDFs, male) were obtained from AllCells, LLC (Alameda, CA), as a pool of three separate donors. Kidney cells were described in [31]. All cells except kindey and MCF10A were maintained in high glucose DMEM (Invitrogene) with 5% FBS (different vendors) and antibiotic solution in standard conditions (5% CO2, 37°C). Composition of the medium for MCF10A cells is provided in Table S1. Kidney cells were maintained as described in [31]. All cells were routinely tested for mycoplasma using Lonza’s MycoAlert® Mycoplasma Detection Assays (Lonza, Rochester, NY).

### Cell transformation

NDF and MCF10A cells were transduced with lentivirus harboring p53 dominant negative mutant GSE56 and HRasV12 oncogene connected via IRES [36]. Control cells were transduced with empty virus. Cells were transduced at MOI ∼ 1. After that cells were split every 3-4 days for 2 weeks and then taken to the experiments.

MEFs were isolated from C57Bl/6 p53-heterozygous bred mice. Isolated MEFs were genotyped for the absence of p53, gender of embryos was not identified, and p53-null MEFs were immortalized by stable transfection with SV40 large T antigen and then transformed with a vector expressing oncogenic mutant HRasV12 tagged with GFP [36]. GFP positive cells were sorted flow cytometry for further experiments as a model of transformed cells. To grow p53-null transformed MEFs (MEF p53KO-HRasV12) *in vivo*, 0.5×10^6^ cells were implanted subcutaneously into wild-type C57BL/6 mice. When tumors reached 500-600mm^3^, they were excised and underwent enzymatic digestion (1 mg/mL collagenase type IV + 0.02 mg/mL DNase) for 1 hour at 37°C. Digested tumors were filtered through 70 μm mesh filter and plated in DMEM supplemented with 10% FBS.

### Cell treatment, fixation, and staining

For all imaging experiments, cells were plated into 96 well black plates with clear bottoms (Greiner Bio-One, Monroe, NC, cat # 655090), 5000 cells per well. For flow cytometry cells were plated in 6 well plate at 100,000 cells per well. Next day cells were treated with CBL0137 for 10-60 min or epigenetic drugs for 24 hours. After treatment medium with drugs was removed and cells were fixed with 4% paraformaldehyde (PFA) in PBS with 0.1% Triton X100 or 100% ice-cold methanol for 10 min at room temperature. For flow cytometry cells were first trypsinized, resuspended in medium with 5% FBS, washed from medium with PBS and fixed in 4% PFA. Then fixatives were removed, and cells were stained with DNA ligands with or without RNAse A (100 μg/ml). Unless otherwise stated, the following concentrations of DNA ligands were used Propidium Iodide (PI) –1 μg/ml, DAPI and Hoechst 33342 – 1 μM, SYBR Green – 50 μM (1:100 dilution of solution provided by vendor). Plate was left in staining solution overnight at room temperature to allow RNA digestion.

### Image acquisition and analyses

Plate imaging was done using Cytation 5 automated imager (Biotek/ Agilent Technologies, Santa Clara, CA). We used a 4X and 10X objectives, 4 images or 9 images per well of 96-well plate, four replicate wells per condition. Data obtained with 4X and 10X objectives were very similar, therefore we decided to do most of the study experiments with 4X objective due to the lower data volume without compromised accuracy. Cytation 5 built-in Gen5 software (Agilent) creates a montage of images taken from the same well. We used ‘autofocus’ mode, which ensures that each image is in focus independently of potential irregularities in plastic thickness. Data collection was done from ∼300-5000 objects per well. Object masking was done to include only nuclei. For the same DNA stain, we set the same exposition time and object masking parameters for the whole experiment. If experiment consisted of several plates, each one of them included positive/negative controls for comparison. This serves as intrinsic normalization for the data collection. The data we collected and included into the manuscript included (i) ‘size’ -the longest linear dimension of an object/nucleus (in μm), (ii) ‘area’ – the square of an object/nucleus (in μm2). These parameters are specific to the camera (CMOS), and the objectives used, and are gained from an image as the number of pixels per object (or per the longest dimension of an object for ‘size’) multiplied by the square (or by length for ‘size’) of each pixel. In our case, for images taken with 4x objective the pixel length was 1.28 μm and the area - 1.64825 μm2. (iii) ‘Mean’ - is the intensity of fluorescence of one pixel of an object, averaged for all object pixels. (iv) ‘Integral’ - is the sum of the fluorescent intensities of all pixels from an object. Before quantitation of fluorescent intensity, objects from all wells were sorted by size, and objects <15 μm and > 40 μm were excluded to remove cell debris or cell clusters. Using this filtered data, violin plots or histograms showing the distribution of signal between different objects within the same experimental condition were built using Seaborn/Matplotlib violinplot or distplot functions. Microscopic imaging was done using Zeiss Axio Observer A1 inverted microscope, Zeiss MRC5 camera, and AxioVision Rel.4.8 software.

### Statistical analyses

All experiments were repeated at least twice and included at least four replicate wells. The average parameter for a well was used to calculate the mean of replicate wells. For object level analyses, all objects from replicate wells were pooled together. The significance of difference between conditions was calculated using methods indicated in figure legends.

## Supplementary figures

**Supplementary Figure S1.**
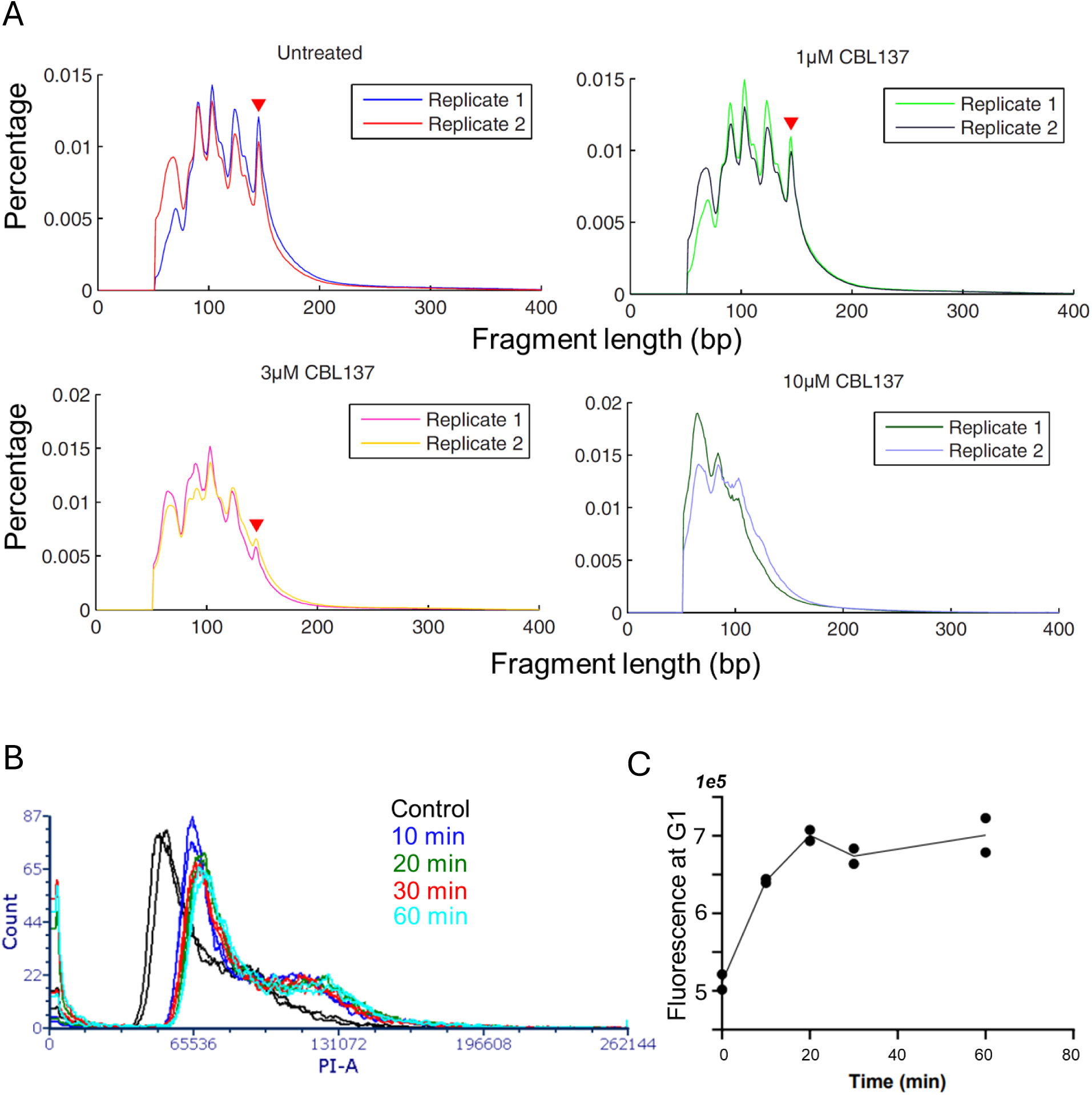
A. Distribution of fragments length obtained from MNAse digested chromatin in HT1080 cells, untreated or treated with different doses of CBL0137 for 1 hour. Two biological replicates are shown. Red triangle indicates the peak corresponding to the size of fully wrapped nucleosomal DNA of 147 bp. This fragment is completely lost in samples treated with 10 μM of CBL0137. B, C. Distribution of PI fluorescence of HT1080 cells treated with 10 μM of CBL0137 for the different amount of time analyzed using flow cytometry. Two replicates per condition were used. B. Histograms of distribution. C. Fluorescent intensity of G1 peak from B.

**Supplementary Figure S2.**
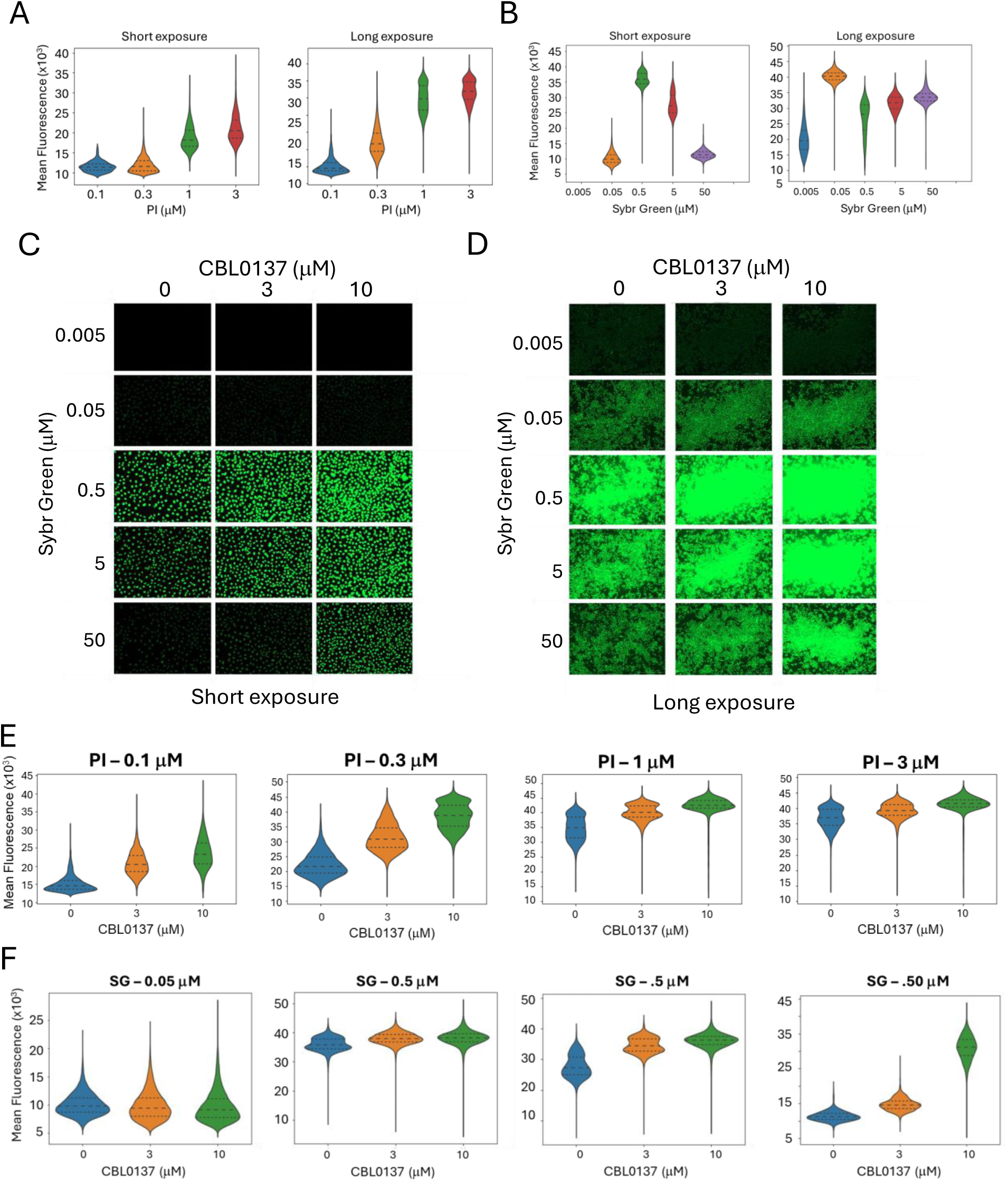
A. Titration of DNA intercalators for optimal measurement of chromatin accessibility. Replicate sets of HT1080 cells, treated for 30 minutes with CBL0137 were fixed and stained with the indicated concentrations of PI (A, E) or Sybr Green (B, C, D, F). Short exposure was selected for optimal performance of the brightest cells. Long exposure was selected for the optimal performance of the dimmest cells. No fluorescence was detected in wells stained with 0.005 μM of Sybr Green. A,B. Violin plots of mean nuclear fluorescence of PI (A) or Sybr Green (B). C, D. Short and long exposure images of wells stained with Sybr Green. Images on panel C are enlarged 10 times comparing with images on panel D. E, F. Violin plots comparing performances of different concentrations of PI (E) or Sybr Green in HT1080 cells treated with different concentrations of CBL0137.

**Supplementary Figure S3.**
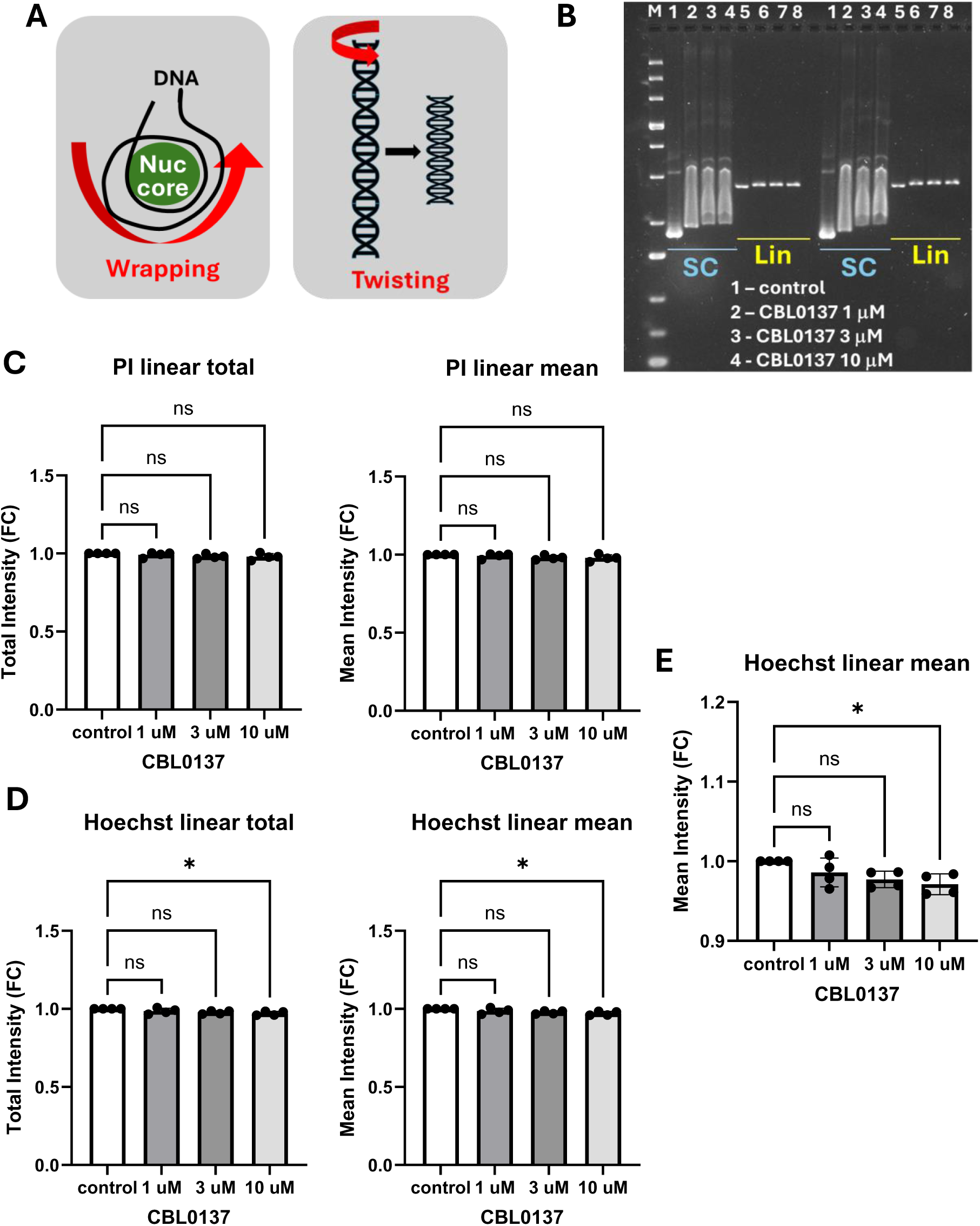
Increase of DNA fluorescence upon CBL0137 treatment is not due to DNA untwisting. A. Schematic presentation of the difference between terms “wrapping”, which means winding of DNA around nucleosome core, and “twisting”, rotation of DNA helix around its axis. B. Example of gel shift experiment in which fluorescence of linear (Lin) DNA bands in the presence of CBL0137 was compared with drug-free linear DNA (control). SC – supercoiled plasmid is shown to demonstrated the shift in DNA velocity due to the presence of the drug. Two technical replicates are shown. C-E. Average band intensity of PI (C) or Hoechst (D, E) presented as fold changes versus control in each experiment. N= 4 (two experiments with two replicates as in B). E. Mean linear fluorescence of Hoechst shown with Y-axis limited to 0.9 – 1.2 to show the reduction of fluorescence of CBL0137 samples. Asterisk – p < 0.05 using one-way ANOVA.

**Supplementary Figure S4.**
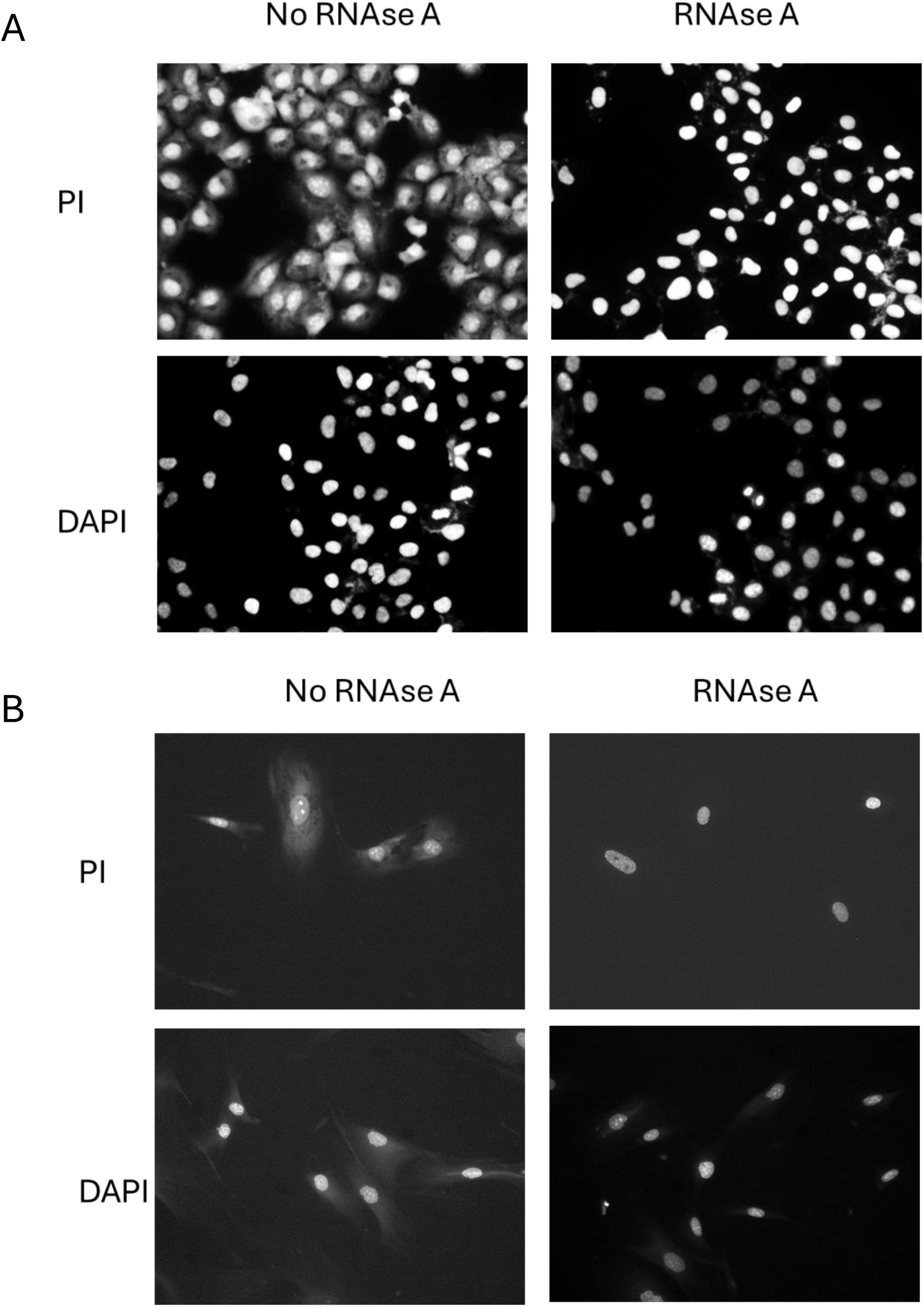
Effect of RNAse A on the staining of HT1080 (A) or NDF (B) cells with PI and DAPI. Microscopic images of cells stained with the indicated dyes in the presence or absence of RNAse A

**Supplementary Figure S5.**
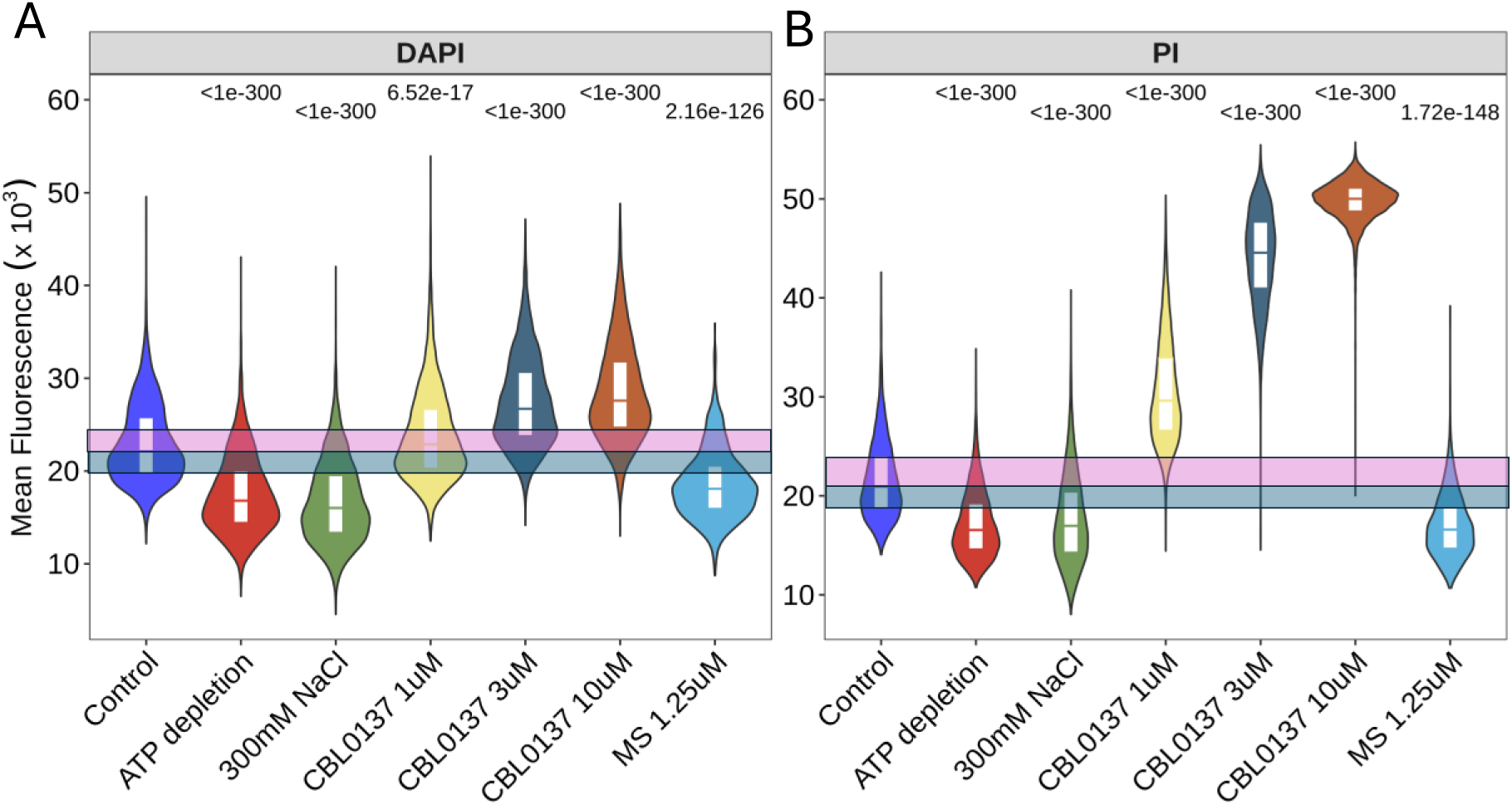
Distribution of mean nuclear fluorescence in HT1080 cells treated with different means leading to chromatin compaction, including ATP depletion, hypertonic shock (300 mM of NaCl), CBL0137 as a control for chromatin decompaction and methylstat, inhibitor of histone demethylases. A. Staining of treated cells with DAPI. B. Staining of replicate cells with PI. Pink and blue transparent squares show positions of quartiles 0.5 and 0.75 (pink), and 0.5 and 0.25 (blue) in control untreated samples. Numbers above violin plots show Holm adjusted p-values for Kruskal-Wallis test with post-hoc Dunn’s test comparing treated cells and control cells.

**Supplementary Figure S6.**
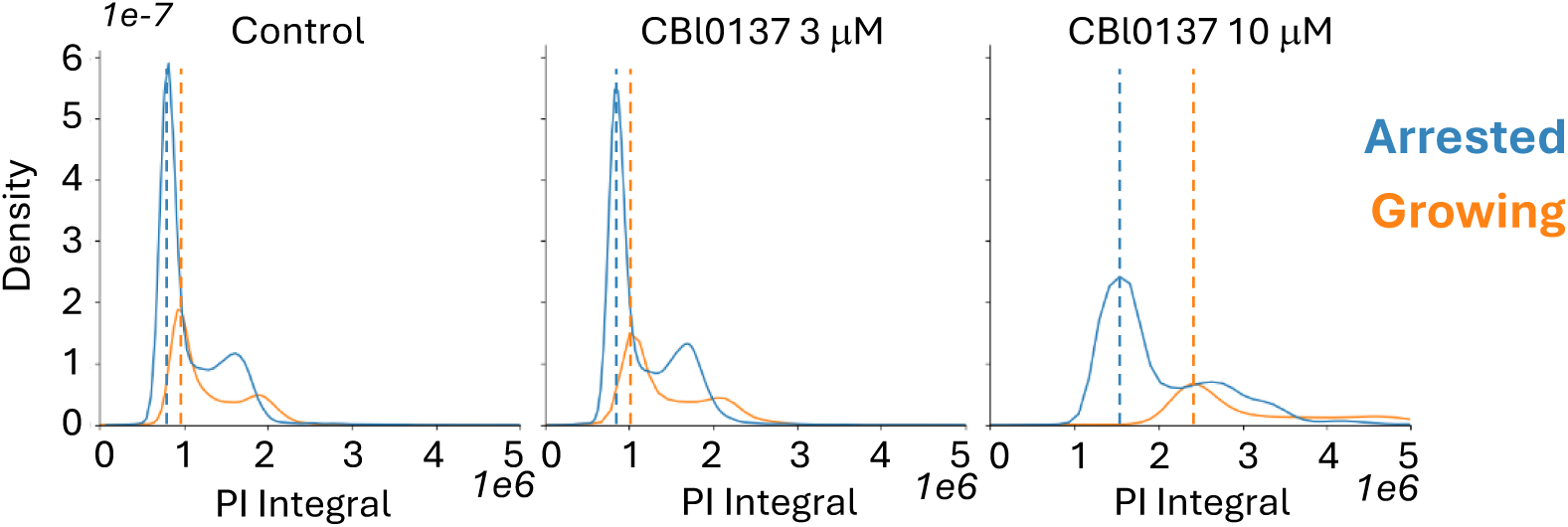
Distribution of total nuclear fluorescence in growing or arrested HT1080 cells treated with CBL0137 and stained with PI. Dotted lines showed the positions of G1 peaks in growing (orange) and arrested (bleu) cells.

**Supplementary Figure S7.**
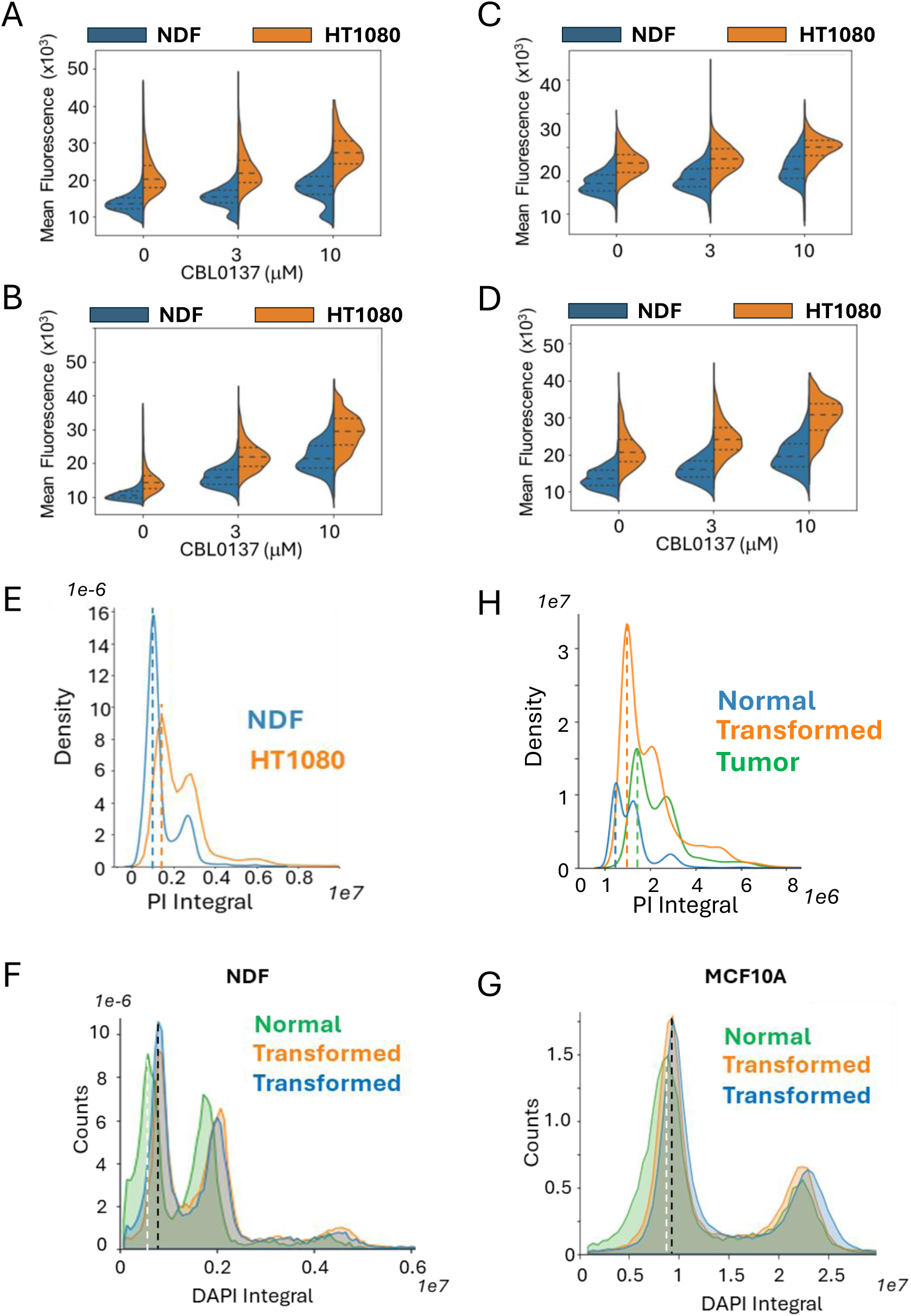
Difference in total nuclear fluorescence between normal, transformed and tumor cells. A-D. Split violin plot with quartiles showing mean fluorescent intensity of NDF and HT1080 cells stained with PI after fixation with PFA (A, B) or methanol (C, D) in the absence (A, C) or presence of RNAse A. Before fixation cells were treated for 30 minutes with the indicated concentrations of CBL0137. E. Comparison of positions of G1 peaks in untreated NDF (blue dotted line) and HT1080 cells (orange dotted line). F, G. Comparison of the positions of G1 peaks in non-transformed NDF (F) or MCF10A (G) cells (white dotted line) and corresponding transformed cells (two biological replicates, black dotted lines). H. Comparison of positions of G1 peaks in normal MEFs (blue dotted like), transformed MEFs (orange dotted line) and tumors established from transformed MEFs (green dotted line). E, H - kernel density estimate (KDE) plots, F, G – histogram plots.

**Supplementary Figure S8.**
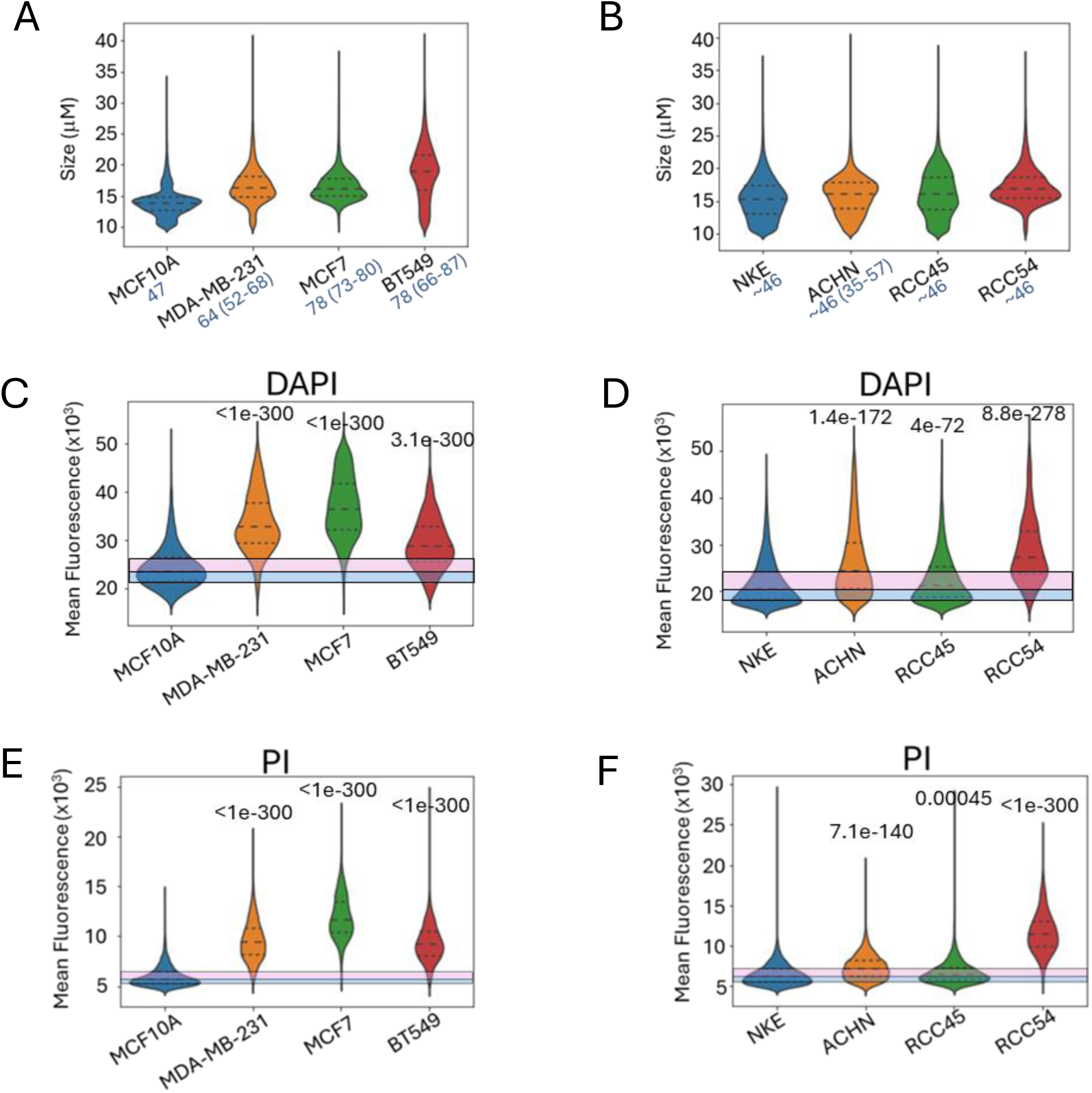
Difference in nuclear size and mean nuclear fluorescence between human normal and tumor cell lines of breast (A, C, E) or kidney (B, D, F). A, B. Violin plot with quartiles showing distribution of nuclear sizes for cell lines. Blue numbers next to the cell line name number of chromosomes reported for these cells are shown. C - F. Violin plots with quartiles showing distribution of mean fluorescent intensity of DAPI (C,D) or PI (E, F) stained cells. Pink and blue transparent squares show positions of quartiles 0.5 and 0.75 (pink), and 0.5 and 0.25 (blue) in control untreated samples. Number above violin is Holm adjusted p-value of for Kruskal-Wallis test with post-hoc Dunn’s test comparing each tumor cell line with non-tumor cell line of corresponding origin, rounded to 3 decimal points.

## Notes

### Competing Interest Statement

The authors have declared no competing interest.

### Summary of Updates

in response to the reviewers comments. Several new experiments were performed and described.

