## Supplementary material for "Image-Based Quantitative Single-Cell Method Suggests Increase of Global Chromatin Accessibility in Tumor Compared to Non-tumor Cell Lines": Fig. S

### Supplementary figures

**Table 1: Media recipes for MCF-10A Cells**

| **Component** | **Growth Medium**[1] | **Resuspension Medium**[1] | **Assay Medium**[1] **(Without EGF)** |
|---|---|---|---|
| DMEM/F12 (Invitrogen #11330-032) | 500.0 ml | 400.0 ml | 500.0 ml |
| Horse Serum (Invitrogen#16050-122) | 25.0 ml (5% final) | 100.0 ml (20% final) | 10.00 ml (2% final) |
| EGF (100µg/ml stock)[2] | 100 µl (20ng/ml final) | -- | -- |
| Hydrocortisone (1mg/ml)[3] | 250 µl (0.5 mg/ml final) | -- | 250 µl (0.5 µg/ml final) |
| Cholera Toxin (1mg/ml stock)[4] | 50 µl (100 ng/ml final) | -- | 50µl (100 ng/ml final) |
| Insulin (10mg/ml stock)[5] | 500 µl (10µg/ml final) | -- | 500µl (10µg/ml final) |
| Pen/Strep (100x solution, Invitrogen #15070-063) | 5.0 ml | 5.0 ml | 5.0 ml |

**Notes:**

1) For each medium type, premix all of the appropriate additives, sterile filter through a 0.2 µm filer, and add to DMEM/F12 medium bottle.
2) EGF: (Peprotech, 1 mg): Resuspend at 100 µg/ml in sterile $dH_2O$. Store aliquots at –20°C.
3) Hydrocortisone: (Sigma #H-0888, 1 g bottles) Resuspend at 1 mg/ml in 200 proof ethanol and store aliquots at -20 °C.
4) Cholera Toxin: (Sigma #C-8052, 2 mg vials) Resuspend at 1 mg/ml in sterile $dH_2O$ and allow to reconstitute for about 10 minutes. Store aliquots at 4°C.
5) Insulin: (Sigma #I-1882, 100 mg vials) Resuspend at 10 mg/ml in sterile $dH_2O$ containing 1% glacial acetic acid. Shake solution and allow 10-15 min to reconstitute. Store aliquots at –20C.

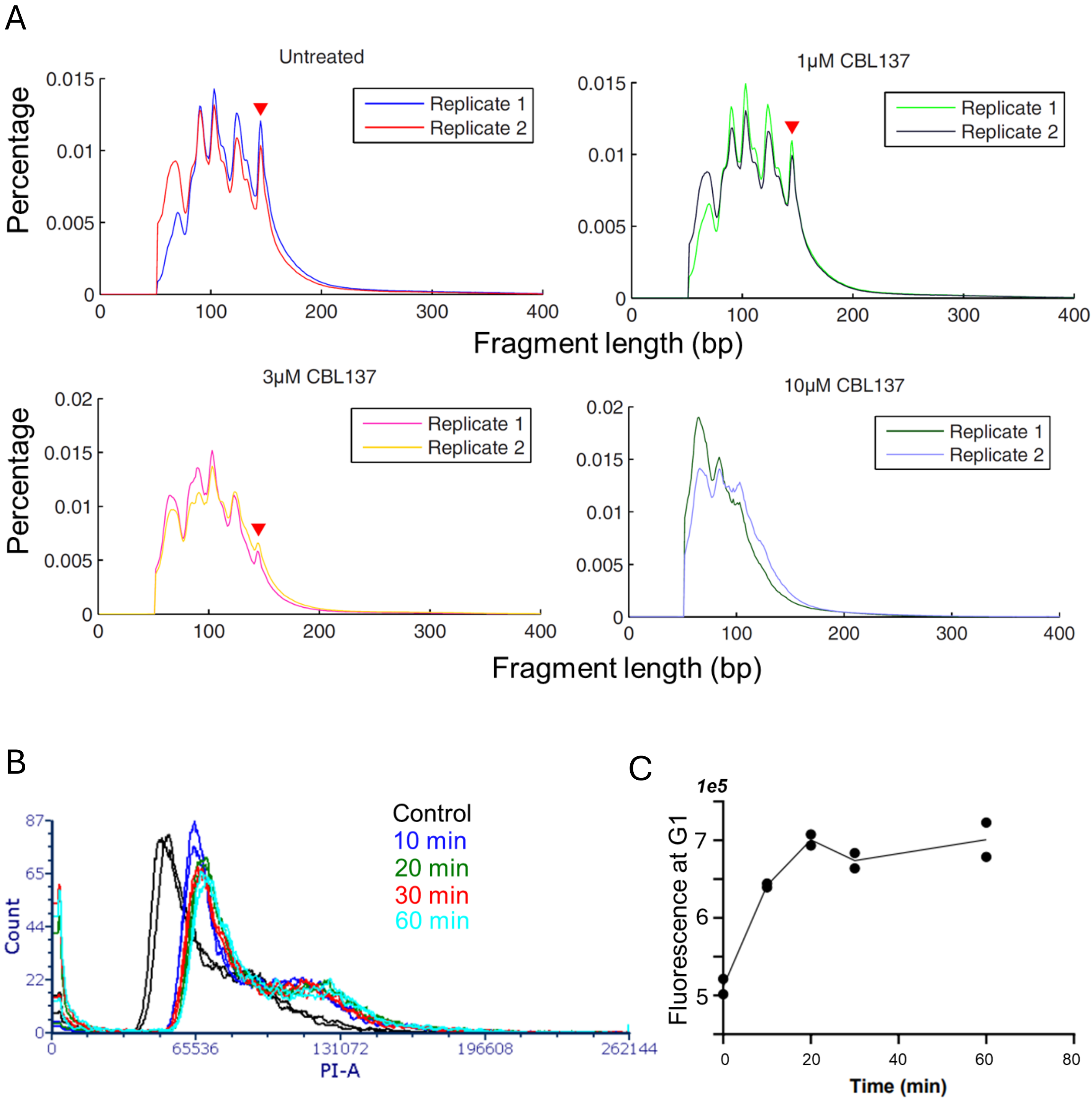


Supplementary Figure S1. A. Distribution of fragments length obtained from MNAse digested chromatin in HT1080 cells, untreated or treated with different doses of CBL0137 for 1 hour. Two biological replicates are shown. Red triangle indicates the peak corresponding to the size of fully wrapped nucleosomal DNA of 147 bp. This fragment is completely lost in samples treated with 10 μM of CBL0137. B, C. Distribution of PI fluorescence of HT1080 cells treated with 10 μM of CBL0137 for the different amount of time analyzed using flow cytometry. Two replicates per condition were used. B. Histograms of distribution. C. Fluorescent intensity of G1 peak from B.

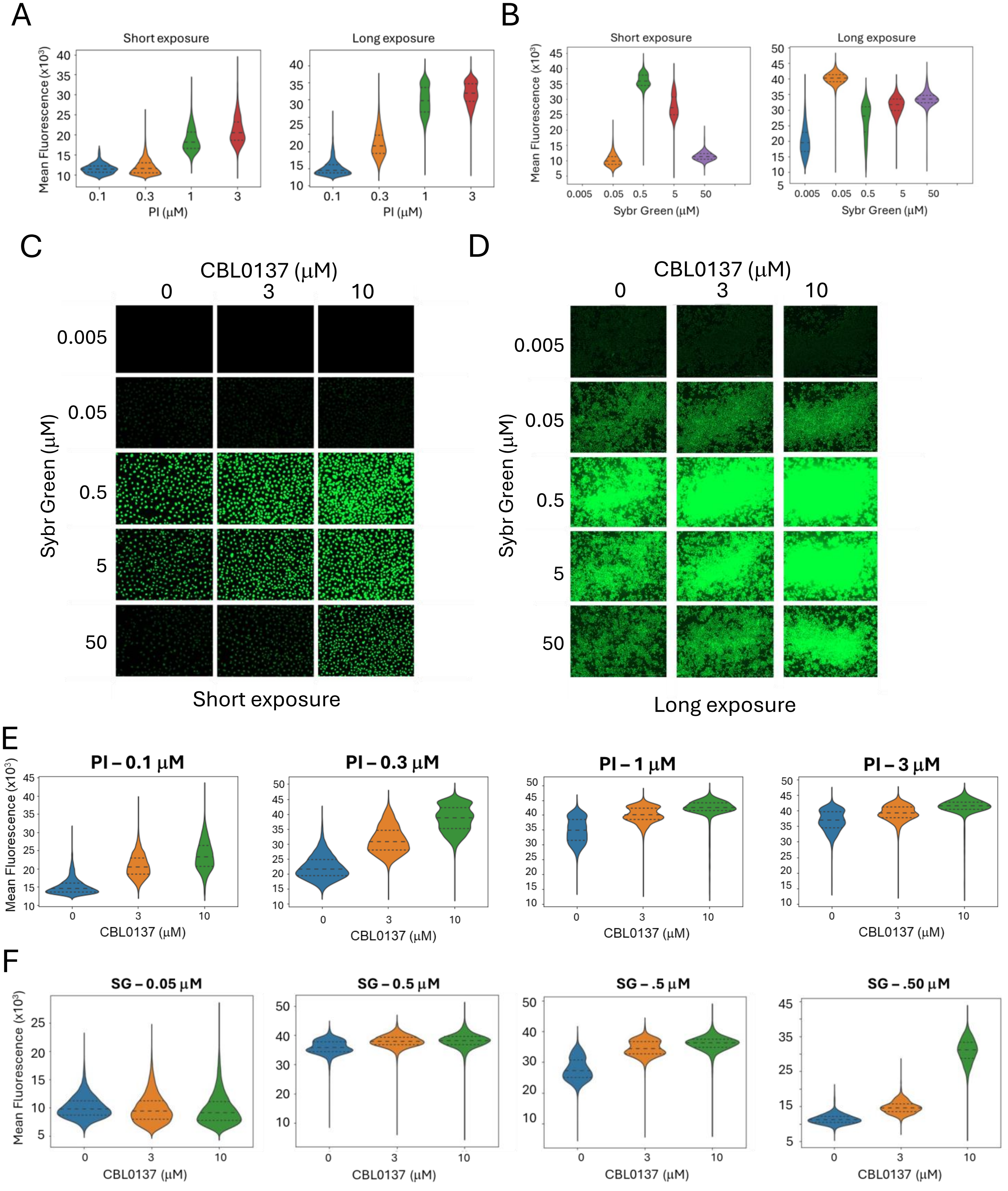


Supplementary Figure S2. A. Titration of DNA intercalators for optimal measurement of chromatin accessibility. Replicate sets of HT1080 cells, treated for 30 minutes with CBL0137 were fixed and stained with the indicated concentrations of PI (A, E) or Sybr Green (B, C, D, F).

Supplementary Figure S2 *continued*.

Short exposure was selected for optimal performance of the brightest cells. Long exposure was selected for the optimal performance of the dimmest cells. No fluorescence was detected in wells stained with 0.005 μM of Sybr Green. A,B. Violin plots of mean nuclear fluorescence of PI (A) or Sybr Green (B). C, D. Short and long exposure images of wells stained with Sybr Green. Images on panel C are enlarged 10 times comparing with images on panel D. E, F. Violin plots comparing performances of different concentrations of PI (E) or Sybr Green in HT1080 cells treated with different concentrations of CBL0137.

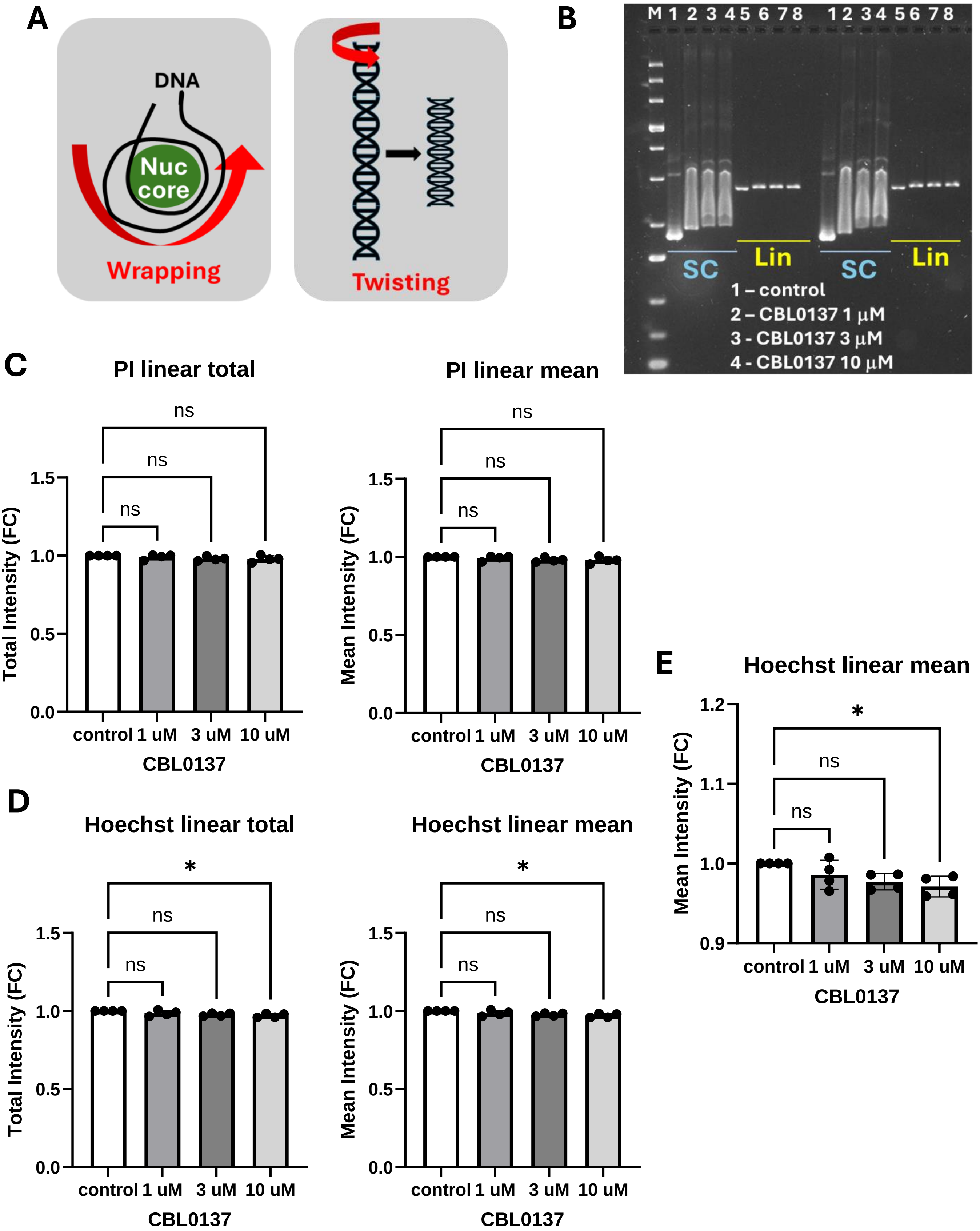


Supplementary Figure S3. Increase of DNA fluorescence upon CBL0137 treatment is not due to DNA untwisting. A. Schematic presentation of the difference between terms “wrapping”, which means winding of DNA around nucleosome core, and “twisting”, rotation of DNA helix around its axis.

Supplementary Figure S3 *continued*.
B. Example of gel shift experiment in which fluorescence of linear (Lin) DNA bands in the presence of CBL0137 was compared with drug-free linear DNA (control). SC – supercoiled plasmid is shown to demonstrated the shift in DNA velocity due to the presence of the drug. Two technical replicates are shown. C-E. Average band intensity of PI (C) or Hoechst (D, E) presented as fold changes versus control in each experiment. N= 4 (two experiments with two replicates as in B). E. Mean linear fluorescence of Hoechst shown with Y-axis limited to 0.9 – 1.2 to show the reduction of fluorescence of CBL0137 samples. Asterisk – $p < 0.05$ using one-way ANOVA.

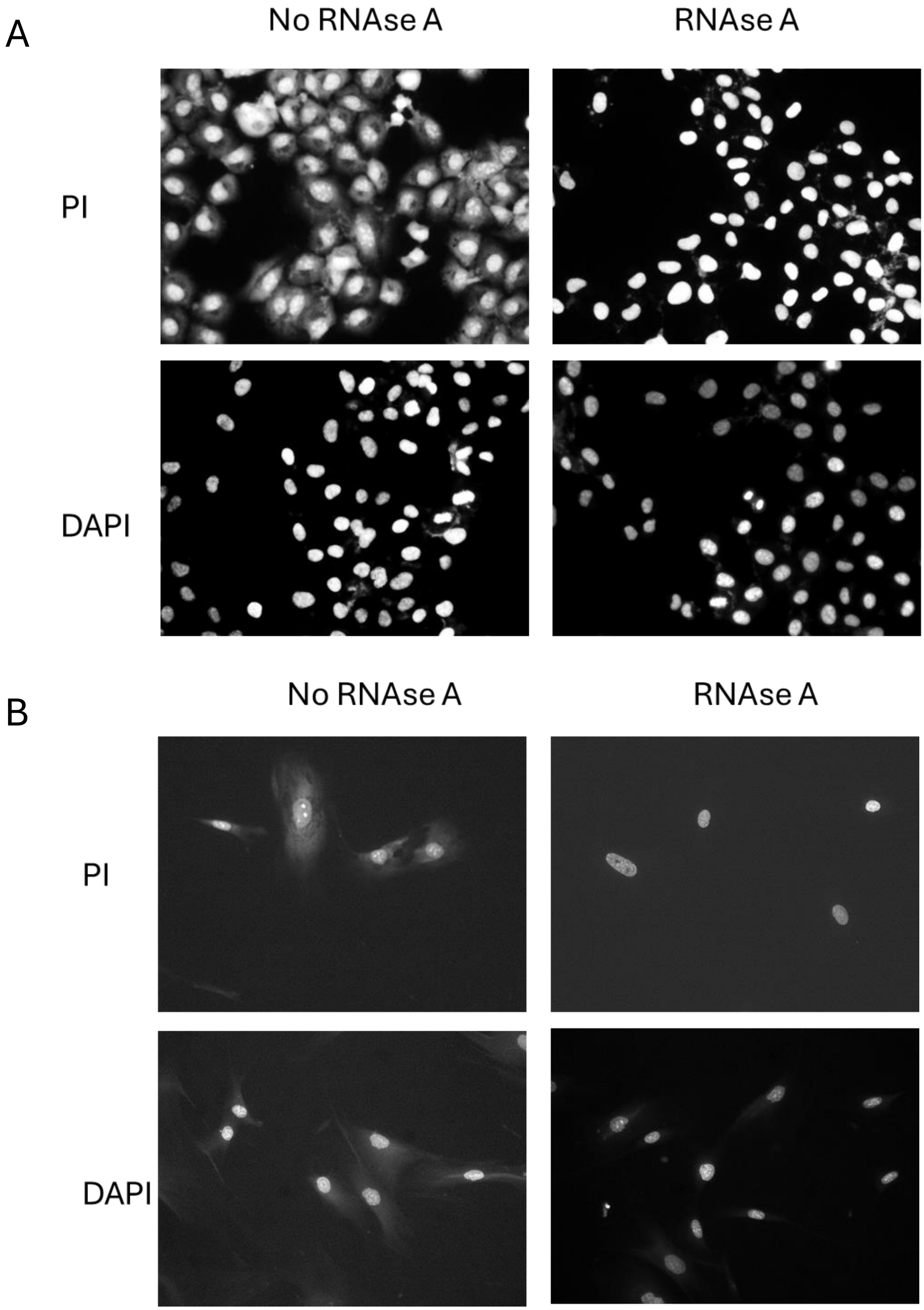


Supplementary Figure S4. Effect of RNAse A on the staining of HT1080 (A) or NDF (B) cells with PI and DAPI. Microscopic images of cells stained with the indicated dyes in the presence or absence of RNAse A

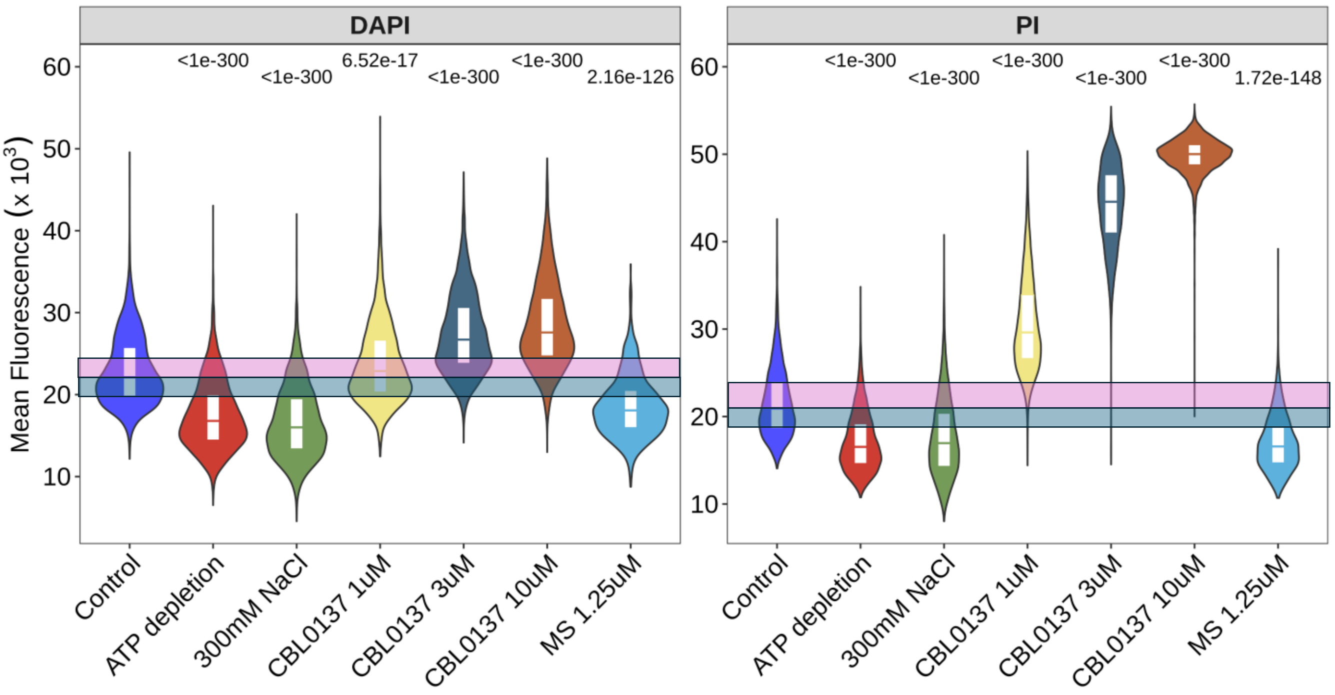


Supplementary Figure S5. Distribution of mean nuclear fluorescence in HT1080 cells treated with different means leading to chromatin compaction, including ATP depletion, hypertonic shock (300 mM of NaCl), CBL0137 as a control for chromatin decompaction and methylstat, inhibitor of histone demethylases. A. Staining of treated cells with DAPI. B. Staining of replicate cells with PI. Pink and blue transparent squares show positions of quartiles 0.5 and 0.75 (pink), and 0.5 and 0.25 (blue) in control untreated samples. Numbers above violin plots show Holm adjusted p-values for Kruskal-Wallis test with post-hoc Dunn’s test comparing treated cells and control cells.

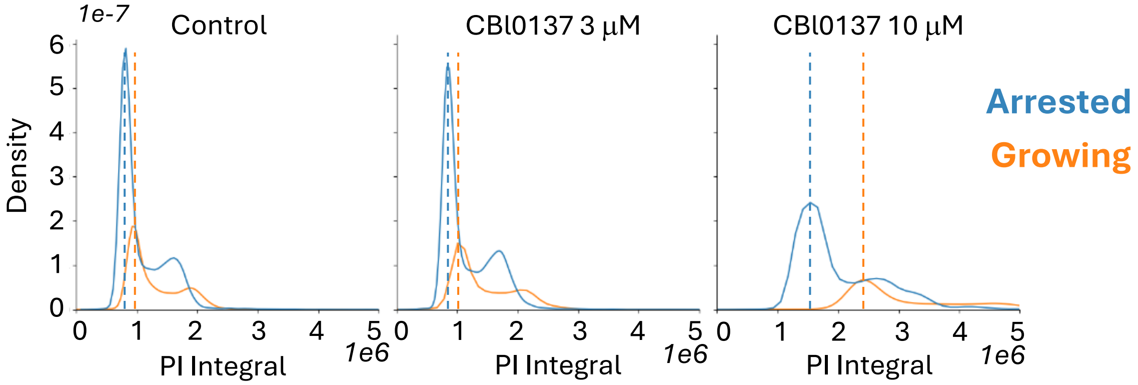


Supplementary Figure S6. Distribution of total nuclear fluorescence in growing or arrested HT1080 cells treated with CBL0137 and stained with PI. Dotted lines showed the positions of G1 peaks in growing (orange) and arrested (bleu) cells.

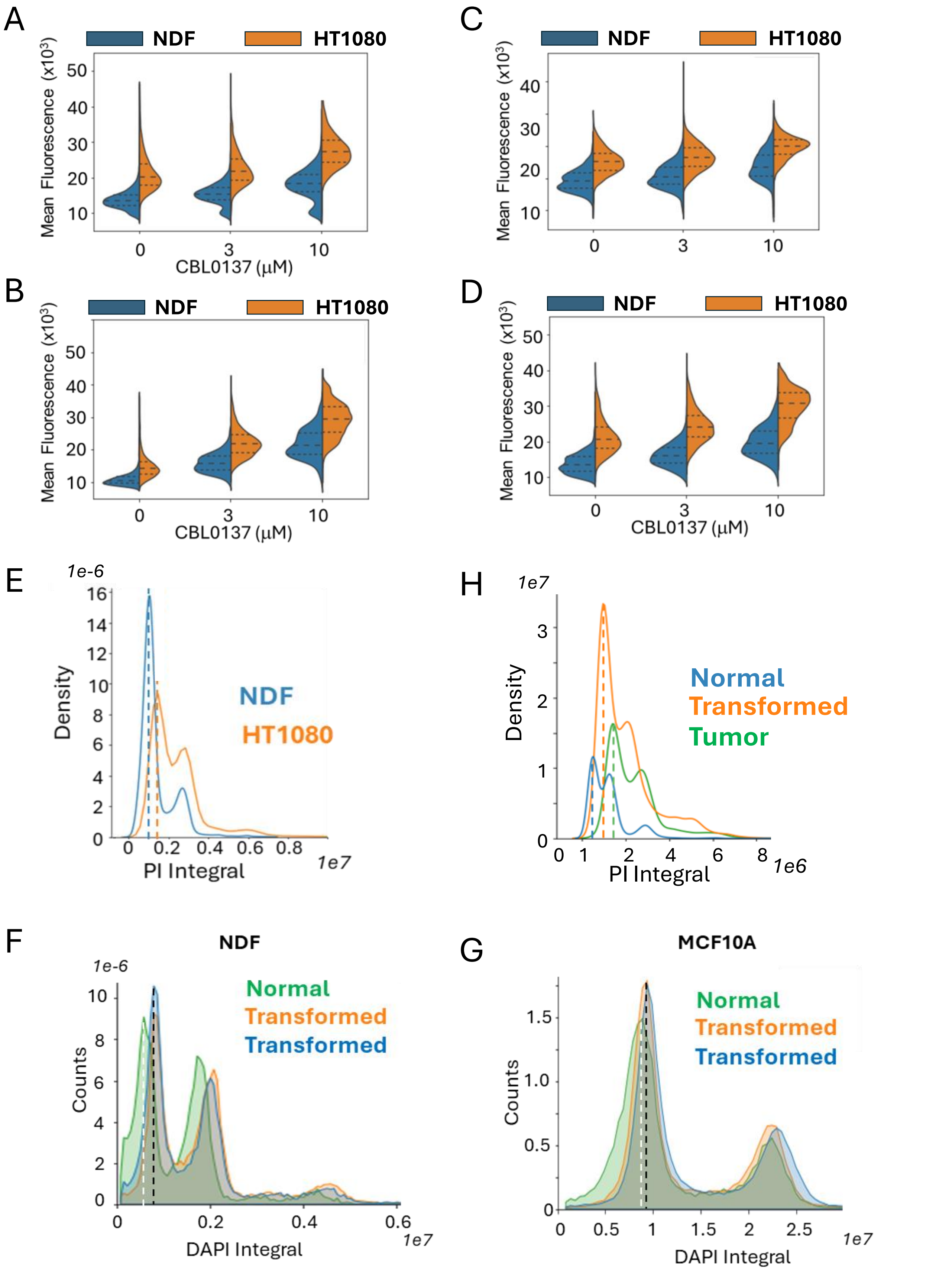


Supplementary Figure S7. Difference in total nuclear fluorescence between normal, transformed and tumor cells.

Supplementary Figure S7. Difference in total nuclear fluorescence between normal, transformed and tumor cells. A-D. Split violin plot with quartiles showing mean fluorescent intensity of NDF and HT1080 cells stained with PI after fixation with PFA (A, B) or methanol (C, D) in the absence (A, C) or presence of RNAse A. Before fixation cells were treated for 30 minutes with the indicated concentrations of CBL0137. E. Comparison of positions of G1 peaks in untreated NDF (blue dotted line) and HT1080 cells (orange dotted line). F, G. Comparison of the positions of G1 peaks in non-transformed NDF (F) or MCF10A (G) cells (white dotted line) and corresponding transformed cells (two biological replicates, black dotted lines). H. Comparison of positions of G1 peaks in normal MEFs (blue dotted like), transformed MEFs (orange dotted line) and tumors established from transformed MEFs (green dotted line). E, H - kernel density estimate (KDE) plots, F, G – histogram plots.

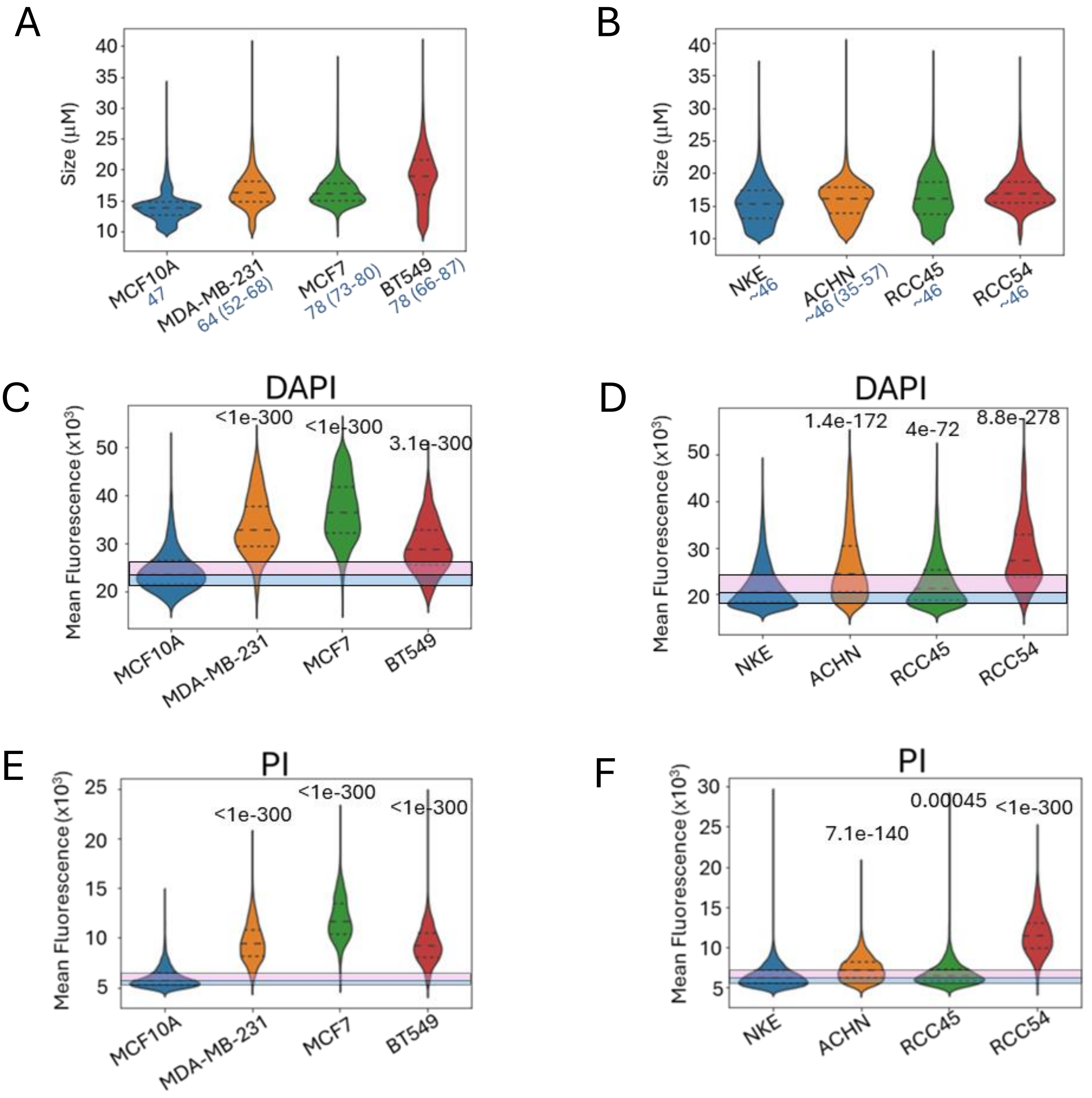


Supplementary Figure S8. Difference in nuclear size and mean nuclear fluorescence between human normal and tumor cell lines of breast (A, C, E) or kidney (B, D, F). A, B. Violin plot with quartiles showing distribution of nuclear sizes for cell lines. Blue numbers next to the cell line name number of chromosomes reported for these cells are shown. C - F. Violin plots with quartiles showing distribution of mean fluorescent intensity of DAPI (C,D) or PI (E, F) stained cells. Pink and blue transparent squares show positions of quartiles 0.5 and 0.75 (pink), and 0.5 and 0.25 (blue) in control untreated samples. Number above violin is Holm adjusted p-value of for Kruskal-Wallis test with post-hoc Dunn's test comparing each tumor cell line with non-tumor cell line of corresponding origin, rounded to 3 decimal points.
